# OSTM1 is a ubiquitin E3 ligase that suppresses B-cell malignancy by activating the cAMP/PKA/CREB pathway

**DOI:** 10.64898/2026.01.23.701155

**Authors:** Muhammad Usama Tariq, Namratha Sheshadri, Jianliang Shen, Jaeyong Jung, Rongrong Li, Kevin Lu, Junrong Yan, Mark C. Koch, Giuseppe Caso, Hassan Sajjad, Brinda Vallat, Stephen K. Burley, Yi Sun, Tong Liu, Hong Li, Christian Hinrichs, Francesco Bertoni, Richard Z. Lin, Jun Wang, Y. Lynn Wang, Jean Vacher, Ping Xie, Wei-Xing Zong

## Abstract

Osteoclastogenesis-associated transmembrane protein 1 (OSTM1) is a membrane-integral glycosylated protein known for regulating lysosomal homeostasis, with loss-of-function mutations causing autosomal recessive osteopetrosis. Through a whole-genome CRISPR/Cas9 screen, we identified OSTM1 as a critical tumor suppressor in B-cell malignancies. In humans, OSTM1 is frequently deleted or downregulated across a wide range of B-cell malignancies. In mice, B-cell-specific monoallelic or biallelic *Ostm1* ablation cooperates with *Cdkn2a* loss to drive lymphomagenesis with near 100% penetrance. Mechanistically, we reveal that a cytosolic, non-glycosylated fraction of OSTM1 functions as an E3 ligase that targets phosphodiesterase 3B (PDE3B) for proteasomal degradation. Because PDE3B catalyzes the conversion of cAMP to AMP and thereby negatively regulating the cAMP-dependent PKA/CREB/CREBBP tumor suppressive pathway, the loss of OSTM1 leads to PDE3B stabilization and enhanced cell transformation. Our findings establish OSTM1 as a pivotal E3 ligase that prevents B-cell lymphomagenesis through the regulation of the cAMP/PKA/CREB pathway.

## Introduction

B-cell lymphoma, leukemia, and myeloma (BCLs) arise from B cells at various developmental stages. Major BCL subtypes include diffuse large B-cell lymphoma (DLBCL), follicular lymphoma (FL), small lymphocytic lymphoma (SLL)/chronic lymphocytic leukemia (CLL), mantle cell lymphoma (MCL), marginal zone lymphoma (MZL), Burkitt lymphoma (BL), B-cell acute lymphoblastic leukemia (B-ALL), hairy cell leukemia (HCL), Waldenström macroglobulinemia (WM), and multiple myeloma (MM).

Genomic alterations such as chromosomal translocations and deletions play essential roles in oncogenesis by activating oncogenes or inactivating tumor suppressor genes (TSGs). A recurrent event in lymphoid and hematological malignancies is the deletion or loss of heterozygosity of the long arm of chromosome 6 (chr. 6q), which has been reported across a wide range of lymphomas and leukemias ^1–5^. Functional studies have previously identified several TSGs on chr. 6q, including PRDM1/BLIMP1, EPHA7, GRIK2, BACH2, CCNC/Cyclin C, TNFAIP3/A20, and FOXO3, whose inactivation could promote lymphoid and myeloid transformation, alone or in cooperation with additional oncogenic events or loss of other TSGs.

Ba/F3 cells are interleukin 3 (IL3)-dependent immortalized yet non-transformed pro-B cells originally isolated from the bone marrow of C3H mice ^6^. Soon after its establishment, the Ba/F3 cell line was used to validate the oncogenic activity of the BCR-ABL tyrosine kinase, the product of the Philadelphia chromosome translocation in chronic myeloid leukemia (CML) ^7^. This study helped launch the wide use of Ba/F3 cells as a tool for studying oncogenesis, as they can be transformed to IL3-independent by oncogenes or inactivation of TSGs ^7–11^. To identify novel TSGs, we performed a genome-wide CRISPR/Cas9 screen to search for genes whose loss confers IL3-independent growth or survival. One such hit encodes OSTM1.

Ba/F3 cells are interleukin 3 (IL3)-dependent immortalized yet non-transformed pro-B cells originally isolated from the bone marrow of C3H mice ^6^. Soon after its establishment, the Ba/F3 cell line was used to validate the oncogenic activity of the BCR-ABL tyrosine kinase, the product of the Philadelphia chromosome translocation in chronic myeloid leukemia (CML) ^7^. This study established Ba/F3 cells as a tool for studying oncogenesis, as they can be transformed to IL3-independent cells by oncogenes or inactivation of TSGs ^7–11^. Using Ba/F3 cells, we performed a genome-wide CRISPR/Cas9 screen to search for novel TSGs, represented by genes whose loss confers IL3-independent growth or survival. One such hit encoded OSTM1, which we demonstrated acts as an E3 ubiquitin ligase that promotes the proteasomal degradation of PDE3B, thereby activating the cAMP-dependent PKA pathway in B lymphocytes and suppressing B-cell lymphomagenesis.

## Results

### Loss of Ostm1 promotes IL3-independent growth in Ba/F3 cells

Because the IL3-dependent Ba/F3 cell line is a classic tool for studying oncogenesis and drug resistance ^7–11^, we performed a genome-wide CRISPR/Cas9 screen to identify genes whose loss can confer IL3-independence. We infected Ba/F3 cells with the mouse GeCKO v2 library and cultured the cells under IL3-deprived conditions. The survivors were subjected to next-generation sequencing to identify enriched single-guide sgRNAs (sgRNAs). This screen enriched more than 1,000 candidate hits, including known TSGs *PTEN, MTAP, TSC2, BRCA2, PTPRK*, and *NF1* (Fig. 1A). We then focused on the hits without previously defined roles in oncogenesis, one of which was OSTM1 (Fig. 1A).

**Figure 1.**
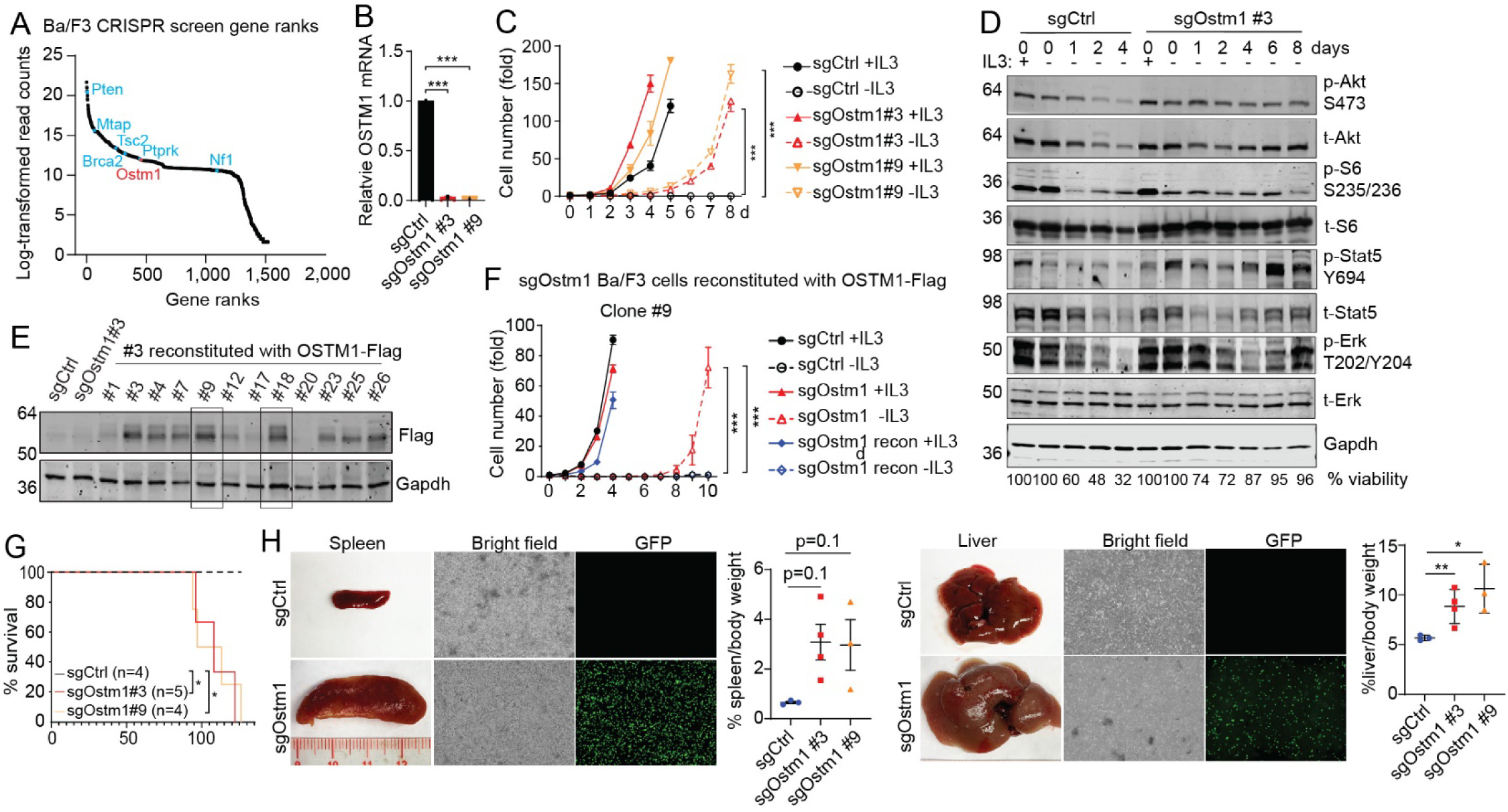
Identification of Ostm1 whose loss promotes IL3-independent growth and transformation of Ba/F3 cells. **(A)** Gene ranking based on log-transformed read counts of enriched hits from a whole-genome CRISPR/Cas9 screen in Ba/F3 cells following IL3 withdrawal. Known TSGs are shown in blue and *Ostm1* in red. **(B)** Ostm1 was silenced using two independent single-guide RNAs (sgRNA) in Ba/F3 cells. Successful silencing was validated by qRT-PCR using on-site primers. **(C)** sgControl and sg*Ostm1* Ba/F3 cell clones were cultured in the presence of absence of IL3. Cell number was measured by Celigo counting and normalized to that of time 0. **(D)** sgControl and sg*Ostm1* Ba/F3 cells were cultured in IL3-deprived medium for the indicated numbers of days. Cell lysates were probed for indicated antibodies. Cell viability was determined by trypan blue exclusion and shown at the bottom of the IB. **(E)** Flag-tagged human OSTM1 (OSTM1-Flag) was reconstituted in sg*Ostm1* Ba/F3 cells. Clones #9 and #18 were selected for further studies. **(F)** sgControl, sg*Ostm1*, and sg*Ostm1* with reconstituted OSTM1-Flag (clone #9) were cultured in the presence or absence of IL3. Cell number was determined. **(G** and **H)** GFP was stably expressed in sgControl and sg*Ostm1* Ba/F3 cell clones. Cells were injected into nude mice (1×10□ cells via i.p. injection). Mice were sacrificed at the endpoints of mice with sg*Ostm1* cells. Kaplan–Meier survival curve was determined **(G)**. Spleens and livers of mice injected with clone #9 cells were photographed and weighed. Splenocytes and hepatocytes were isolated and photographed under bright-field and GFP-fluorescence microscopy **(H)**. Statistical analysis of panels **C** and **F** was done using Student’s t-test, one per row. Survival probability significance for panel **G** was determined using Log-rank (Mantel-cox) test. For bar charts in panels **B** and **H**, one-way ANOVA was used. *p<0.05; **p<0.01; ***p<0.001.

OSTM1 contains 334 amino acids in human (338 in mouse) with a predicted molecular weight of 37 kDa and is ubiquitously expressed (Human Protein Atlas). It was initially discovered in grey-lethal (*gl*) mice, an osteopetrotic mutant that carries a deletion of the *Ostm1* promoter and exon 1 ^12^. In human, loss-of-function mutations in *OSTM1* are associated with autosomal recessive osteopetrosis (ARO), a rare and often fatal skeletal disorder frequently accompanied by neuronal defects ^12–14^. OSTM1 has also been reported to contain a RING-finger motif and to function as a putative ubiquitin E3 ligase (termed GIPN), based on its ability to interact with and promote proteasomal degradation of the Gαi3 subunit of the G protein-coupled receptor (GPCR) complex ^15^. However, the E3 ligase activity of OSTM1 is under debate, and its physiological substrates and biological functions have not been clearly defined.

We verified the effect of *Ostm1* silencing in Ba/F3 cells with two independent sgRNAs distinct from those included in the GeCKO library for our initial screen. Because reliable antibodies for endogenous OSTM1 were then unavailable ^16, 17^, successful *Ostm1* silencing was confirmed by genomic sequencing (Suppl. Fig. S1A) and qRT-PCR (Fig. 1B). *Ostm1* silencing led to IL3-independent growth (Fig. 1C), survival (Fig. 1D), and sustained activation of growth signaling phospho-proteins (Fig. 1D). Reconstitution of human OSTM1 in *Ostm1*-silenced Ba/F3 clones reversed the IL3-independent growth (Fig. 1E and 1F, Suppl. Fig. S1B). To determine whether *Ostm1* silencing promotes tumorigenesis *in vivo*, we stably expressed GFP in sgControl cells and the two independent sg*Ostm1* clones (Fig. 1B) and grafted them into immunodeficient athymic nude mice via tail vein injection. As expected, sgControl-transduced Ba/F3 cells did not form tumors *in vivo*. In striking contrast, both sg*Ostm1* clones resulted in a shortened lifespan (Fig. 1G) and enlarged spleens and livers, characterized by massive infiltration of GFP-expressing cells (Fig. 1H). Therefore, through a whole-genome CRISPR/Cas9 screen, we identified *Ostm1,* whose silencing drives IL3-independent growth and in vivo transformation of Ba/F3 cells. Corroborating the Ba/F3 result, in human BCL lines ARH-77 and SU-DHL-5, silencing OSTM1 led to increased cell proliferation (Suppl. Fig. S1C-S1E). Together, these results suggest that OSTM1 functions as a TSG.

### OSTM1 deletion or down-regulation is associated with human BCL

We then examined OSTM1 genomic alterations in the TCGA database. Among the available cancer types, *OSTM1* deep deletion occurs mostly in DLBCL, prostate cancer, and uveal melanoma (Fig. 2A). Similar to the ARO disease ^12^, no hot-spot mutations were identified in *OSTM1* in BCL patients. Close examination of the literature and publicly available datasets further demonstrated that *OSTM1* deletion occurs across multiple BCL subtypes (Fig. 2B). In the TCGA Mature B-cell Malignancies dataset (MD Anderson Cancer Center), *OSTM1* deletion was observed in several mature B-cell malignancies (Fig. 2C) and was associated with increased genomic instability and poorer survival (Fig. 2D). Of note, in the TCGA Firehose Legacy DLBCL cohort ^18–24^, OSTM1 shallow deletion was observed in 25% and deep deletion in 12.5% of the 48 cases (Suppl. Fig. S2A), which correspond to decreased mRNA level of *OSTM1* (Suppl. Fig. S2B) as well as increased genomic instability (Fig. 2E). At the mRNA level, decreased expression of OSTM1 was observed in multiple BCL patient datasets (Fig. 2F) as well as across BCL cell lines (Fig. 2G). Immunoblotting (IB) revealed that while ectopically expressed OSTM1 had an apparent molecular weight of ∼60 kDa, endogenous OSTM1 ran predominantly at ∼37 kDa in primary healthy peripheral blood mononuclear cells (PBMC), whereas with lower levels or at various ratios between the ∼37 kDa to ∼60 kDa forms in BCL cell lines (Suppl. Fig. S2C and S2D). The ∼37 kDa and ∼60 kDa species were consistent with the previously reported non-glycosylated and glycosylated forms of OSTM1 ^25^. Furthermore, patients with low OSTM1 expression exhibited poorer outcomes across several studies (Fig. 2H). Together, these data indicate that OSTM1 is frequently deleted or downregulated in human BCL, and that its decreased expression correlates with worse prognosis.

**Figure 2.**
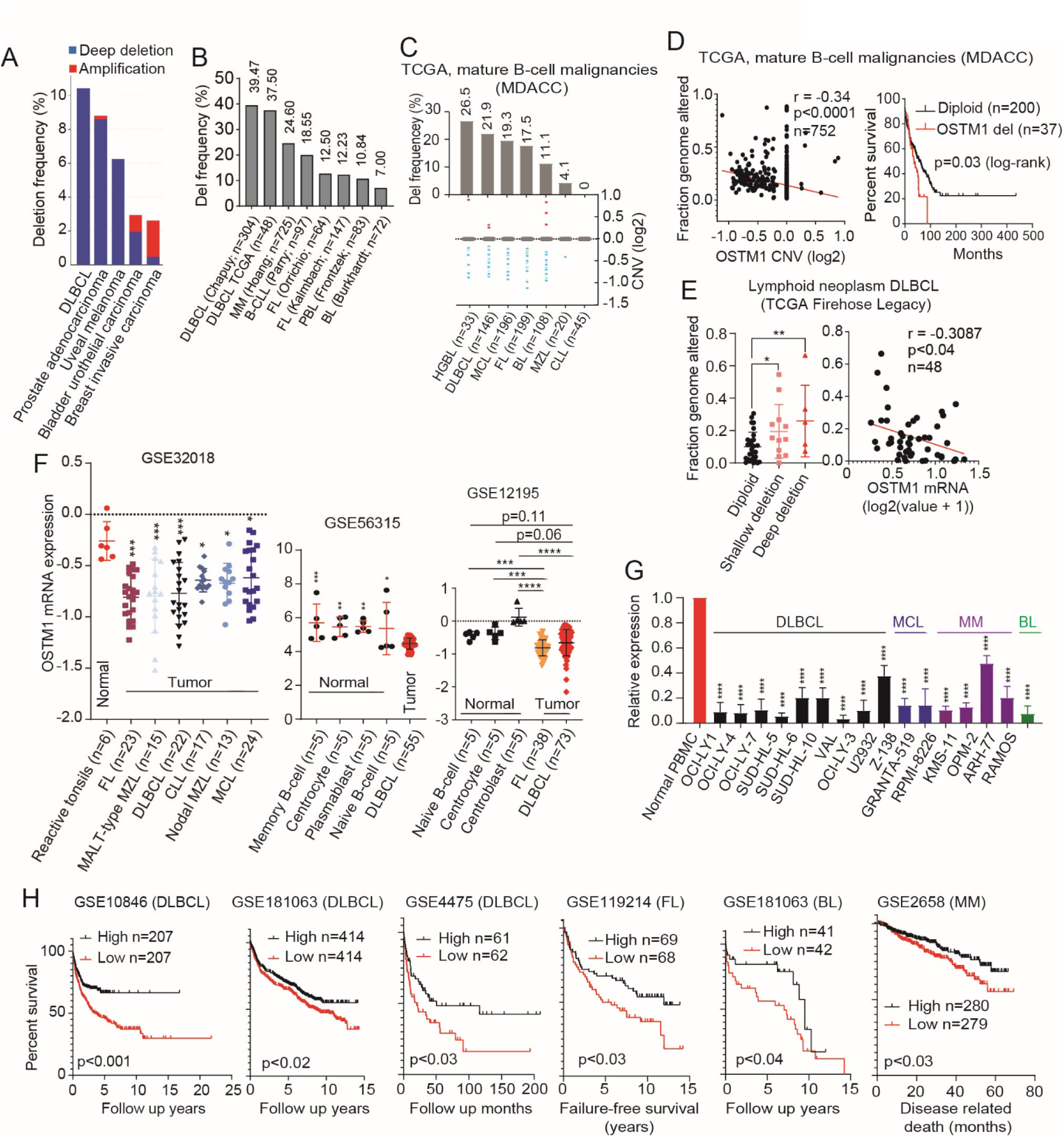
OSTM1 is frequently deleted and down-regulated in human BCL. **(A)** Genomic alterations of *OSTM1* across human cancers were derived from TCGA. Shown are the top 5 cancer types with the most frequent alterations. **(B)** Frequency of *OSTM1* genomic deletions across BCL subtypes from published datasets, including DLBCL, multiple myeloma (MM), follicular lymphoma (FL), B-CLL, and Burkitt lymphoma (BL). **(C)** *OSTM1* log2 copy number variations (CNVs) and percent loss across BCL subtypes in the TCGA Mature B-cell Malignancies dataset (MD Anderson Cancer Center, MDACC). **(D)** Spearman correlations between fraction genome altered (FGA; genomic instability) and *OSTM1* copy number (left) and the Kaplan–Meier survival of patients with or without *OSTM1* deletion in the same dataset in mature B-cell malignancies (dataset from C). **(E)** Correlations between FGA and *OSTM1* deletions (left) or mRNA expression levels (right) in DLBCL patients (TCGA Firehose Legacy). **(F)** *OSTM1* mRNA expression in normal lymphoid tissues vs. BCL subtypes using published datasets. **(G)** qRT-PCR analysis of *OSTM1* expression in cell lines from DLBCL, MCL, MM, and BL compared to PBMCs from healthy donors. **(H)** Kaplan–Meier survival analyses across multiple BCL datasets, stratified by median *OSTM1* expression (upper median: high expression; lower median: low expression).

### Ablation of *Ostm1* cooperates with *Cdkn2a* loss to promote lymphomagenesis

To study the physiological role of OSTM1 in B cells, we crossed *Ostm1^flox/flox^*mice ^26^ with *Cd19*^+/Cre^ mice, in which Cre is expressed in B-lineage cells (from pro-B to mature B cells) ^27^. The resulting B cell-specific *Ostm1* single knockout (SKO) mice were viable and fertile, and no overt disease phenotype was observed. To explore whether *OSTM1* deletion may promote tumorigenesis in cooperation with additional genetic alterations, we searched the TCGA datasets and published studies of human BCLs harboring *OSTM1* deletion for co-occurring alterations in known B-cell TSGs or oncogenes. We found that CDKN2A (located on chr. 9p21) is frequently deleted across a wide spectrum of BCLs, and that OSTM1 deletion frequently co-occurs with CDKN2A deletion (Suppl. Fig. S3A and S3B). Notably, patients with co-deletion of chr. 6q and 9p21 exhibited poorer overall survival (Suppl. Fig. S3C). We thus hypothesized that OSTM1 and CDKN2A act in a cooperative manner to suppress B-cell lymphomagenesis.

To test this hypothesis, we bred *Ostm1^flox/flox^Cd19*^+/Cre^ mice with *Cdkn2a^flox/flox^* mice and generated accumulated cohorts of mice with B cell-specific *Ostm1*^-/-^ (O^-/-^), *Cdkn2a*^-/-^ (C^-/-^), *Ostm1*^+/-^;*Cdkn2a*^-/-^ (O^+/-^C^-/-^), and *Ostm1*^-/-^;*Cdkn2a*^-/-^ (DKO). While O^-/-^ mice did not develop tumors throughout their lifespan, C^-/-^ mice displayed a low penetrance of B-cell lymphoma (Fig. 3A), in agreement with a previous study ^28^. Strikingly, both O^+/-^C^-/-^ and DKO mice exhibited markedly shortened survival (Fig. 3A), accompanied by splenomegaly, lymphadenopathy, and hepatomegaly, as well as spontaneous BCL development, which was detectable in DKO mice as early as 27 weeks of age (Fig. 3B and 3C). Hematoxylin and eosin (H&E) staining and immunohistochemistry (IHC) revealed severe disruption of splenic architecture, with loss of normal red pulp and white pulp boundaries and massive infiltration of proliferating (Ki-67^+^) B220^+^ B cells extending into both the T-cell zones and the red pulp in O^+/-^C^-/-^ and DKO mice (Fig. 3C). Flow cytometry analyses further confirmed a high incidence of BCL in the spleen, lymph nodes (LNs), and bone marrow (BM) of O^+/-^C^-/-^ and DKO mice (Fig. 3D), accompanied by a reduction in normal follicular (FO) and marginal zone (MZ) B cells and an increase in CD21^-^CD23^-^ (double negative, DN) immature or activated B cells (Fig. 3E). B-cell receptor (BCR) profiling revealed that splenic BCLs arising in O^+/-^C^-/-^ mice frequently originated from oligoclonal expansion of malignant B cells, whereas those from DKO mice were predominantly monoclonal (Suppl. Fig. S3D and S3E). All analyzed malignant B-cell clones from both O^+/-^C^-/-^ and DKO mice were IgM^+^ with the exception of one clone with a nonproductive VDJ rearrangement, and none showed evidence of Ig isotype switching or somatic hypermutation (SHM) (Suppl. Fig. S3F). These findings indicate that these BCLs originate from immature B cells or mature pre-germinal center (pre-GC) B cells. Thus, consistent with the clinical evidence that OSTM1 shallow deletions are common in BCL patients (Suppl. Fig. S2 and S3A), our observation that O^+/-^C^-/-^ mice developed a robust B-cell lymphoma phenotype indicates that Ostm1 functions as a tumor suppressor in a haploinsufficient manner in the B-cell lineage.

**Figure 3.**
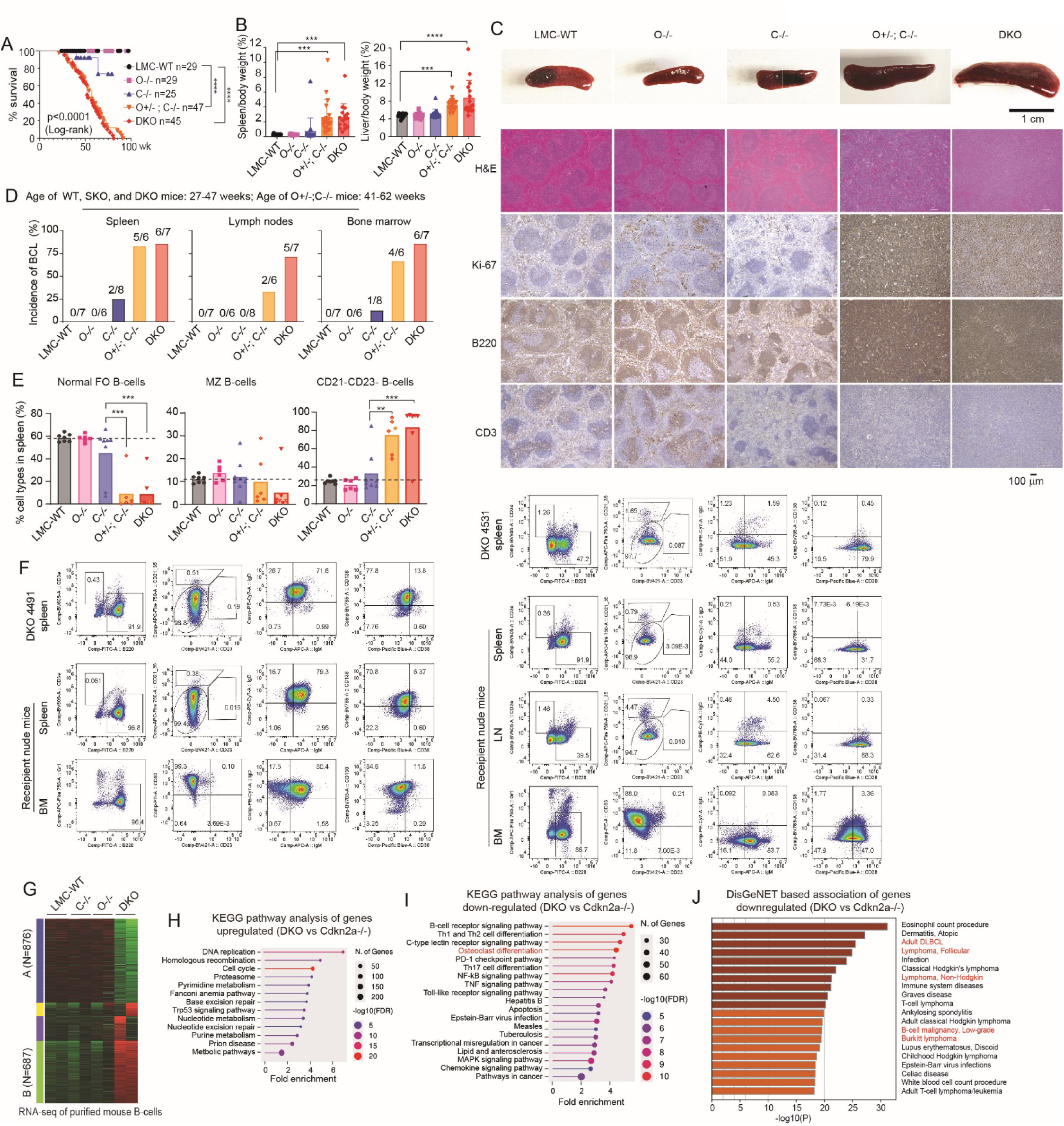
Mono-and biallelic deletion of Ostm1 cooperates with Cdkn2a ablation to promote lymphomagenesis in mice. *Ostm1*^lox/lox^ mice were bred to *Cd19*-Cre and *Cdkn2a*^lox/lox^ mice. **A**) Kaplan-Meier surivival survival curves of littermate control wild-type (LMC-WT), *Ostm1*^-/-^ (O^-/-^), *Cdkn2a*^-/-^ (C^-/-^), *Ostm1*^+/-^;*Cdkn2a*^-/-^ (O^+/-^;C^-/-^), and *Ostm1*^-/-^;*Cdkn2a*^-/-^ (DKO) mice. (**B**) Spleen and liver weight at the time of end-point O^+/-^;C^-/-^ or DKO mice, comparing with the age-and gender-matched WT and single knockout (SKO) mice. (**C**) Spleen images and histopathological features of indicated genotypes analyzed by H&E and IHC staining. (**D**) Incidence of BCL in the spleen, lymph nodes (LNs), and bone marrow (BM), based on clonal abnormal B-cell populations. (**E**) The percentage of normal follicular B cells (FO; B220^+^CD3^-^CD23^+^CD21^int^), marginal zone B cells (MZ; B220^+^CD3^-^CD23^int^CD21^+^), and double negative B cells (B220^+^CD3^-^CD21^-^CD23^-^; activated/immature/malignant B cells) in the spleen of the mice analyzed in (A). **p<0.01; ***p<0.001 as determined by ANOVA. **(F)** Representative transplantation of DKO splenic BCL into nude recipient mice. Splenocytes (1×10^6^) isolated from DKO mice (ID: 4491 or 4531) were injected intraperitoneally (i.p.) into each nude recipient mouse. At 7-9 weeks post-transplantation, the spleen, LN, and BM cells were harvested and analyzed by flow cytometry (FACS). FACS profiles of primary DKO splenocytes are shown for comparison. **(G-J)** Mice of the indicated genotypes were harvested at the endpoint of the DKO cohort (around 31-week-old). Bulk RNA-seq was performed on purified splenic B-cells. **(G)** Heatmap showing genes that were markedly down-regulated (Cluster A) and upregulated (Cluster B) in DKO mice. **(H** and **I)** KEGG pathway enrichment analysis of genes in Cluster A **(H)** and Cluster B **(I)** was performed. The osteoclast differentiation pathway is highlighted. **(J)** Disease-associated gene enrichment analysis of Cluster A (down-regulated genes in DKO mice). B-cell lymphomagenesis-related pathways are highlighted in red.

To further evaluate the malignant potential of the primary splenic BCLs spontaneously developed in DKO mice, we performed transplantation experiments in which mixed splenocytes from individual DKO donor mice were injected intraperitoneally (i.p.) into immunodeficient athymic nude mice. By 7-9 weeks post-transplantation, all recipients developed disseminated lymphoma involving multiple organs, including the spleen, LNs, and BM. The secondary BCLs arising in transplanted nude mice exhibited immunophenotypic features closely resembling those of the spontaneous primary BCLs in DKO donors (Fig. 3F), confirming their malignant and transplantable nature.

To gain transcriptional insights, we performed bulk RNA-sequencing (RNA-seq) on splenic B cells purified from four genotypes (WT, C^-/-^, O^-/-^, and DKO) harvested at the endpoint of the DKO cohort. The most drastic changes were observed in the DKO mice (Fig. 3G). Since Cdkn2a SKO and the WT mice showed similar transcriptome profiles, we compared the DKO with Cdkn2a SKO transcriptomes in detail. Kyoto Encyclopedia of Genes and Genomes (KEGG) pathway enrichment analysis revealed that upregulated genes in the DKO mice were enriched in pathways related to cell growth and proliferation such as DNA replication, repair, cell cycle, and nucleotide synthesis, consistent with the lymphomagenesis phenotype (Fig. 3H). The down-regulated genes were related to osteoclast differentiation as expected, and also to BCR signaling, immune and inflammatory responses (Fig. 3I). DisGeNET analysis further revealed that the down-regulated genes in the DKO were strongly associated with B-cell lymphomagenesis (Fig. 3J). Together, our findings demonstrate that *Ostm1* ablation cooperates with *Cdkn2a* deletion to drive B-cell tumorigenesis.

### OSTM1 interacts with PDE3B and promotes its proteasomal degradation

We next explored the mechanisms by which OSTM1 regulates oncogenesis. We first performed Tandem Mass Tag (TMT)-based mass spectrometry (MS) to identify proteins whose abundance was altered in sg*Ostm1* Ba/F3 cells (Fig. 4A). Notably, proteins identified by TMT-MS included those enriched in the B-cell receptor (BCR) signaling pathway and osteoclast differentiation (Fig. 4B), consistent with the known role of OSTM1 in osteoclast maturation and suggesting its potential role in B-cell biology. We also performed BioID mass spectrometry (BioID-MS) using OSTM1 fused to the biotin ligase BirA to identify OSTM1-interacting proteins in Ba/F3 cells (Suppl. Fig. S4A), in the presence or absence of IL3 (Suppl. Fig. S4B). Comparison of candidates identified by TMT-MS and BioID-MS revealed a single overlapping hit, Pde3b, in Ba/F3 cells cultured under both IL3 conditions (Fig. 4C).

**Figure 4.**
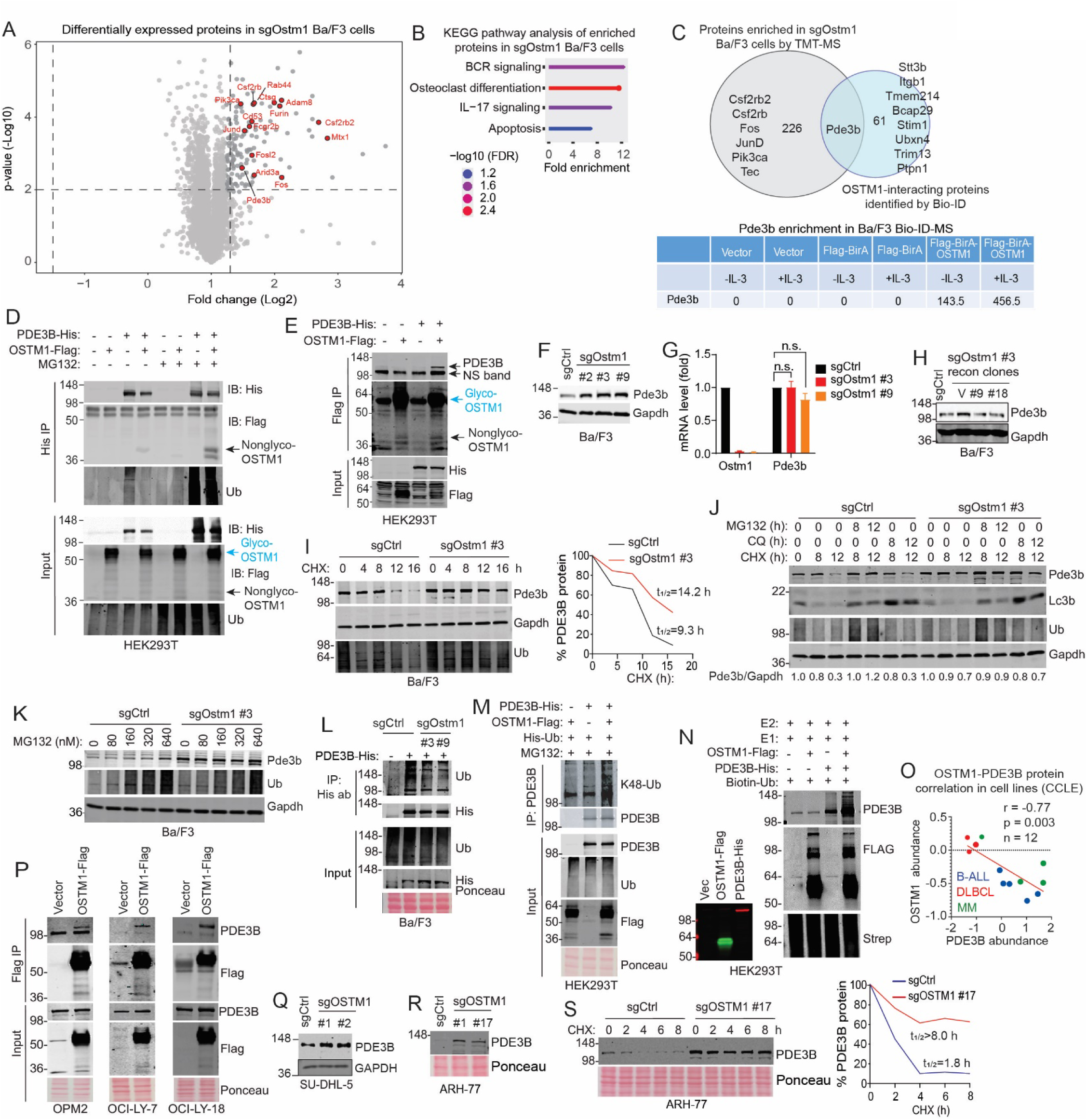
OSTM1 interacts with and ubiquitylates PDE3B to promote its proteasomal degradation. **(A)** sg*Ostm1* and sgControl Ba/F3 cells were subjected to TMT-MS. Volcano plot (log₂ fold change vs. –log₁₀ p-value) was generated using VolcaNoseR. Dashed lines indicate significance cutoffs. Selected proteins involved in hematopoeitic function and differentiation are highlighted in red. **(B)** KEGG pathway enrichment analysis of proteins involved in sg*Ostm1* Ba/F3 cells with-log10(FDR) used as the significance metric. **(C)** Venn diagram depicting the overlap between proteins enriched in sg*Ostm1* cells identified by TMT-MS and OSTM1-interacting partners identified by BioID in Ba/F3 cells. The table shows BioID enrichment scores for Pde3b. **(D and E)** His-tagged PDE3B and Flag-tagged OSTM1 were co-transfected into HEK293T cells, in the presence or absence of MG132 (10 µM). Co-immunoprecipitation (Co-IP) was performed using anti-His (D) or anti-Flag (E) antibodies, followed by IB. **(F)** *Ostm1* was silenced in Ba/F3 cells, and three independent cell clones were probed for Pde3b by IB. **(G)** qRT-PCR analysis of sg*Ostm1* cell clones showing that Pde3b mRNA levels were not affected by *Ostm1* silencing. **(H)** OSTM1-Flag was stably reconstituted in sg*Ostm1* Ba/F3 cell clone #3, and Pde3b was probed by IB. **(I)** sgControl and sg*Ostm1* Ba/F3 cells were treated with cycloheximide (CHX, 100 µg/ml) for the indicated hours. Pde3b was probled by IB and quantified using ImageJ. Pde3b half-life was calculated based on densitometry. Total ubiquitylated proteins were probed as an indicator of global protein turnover following CHX tratment. **(J)** sgControl and sg*Ostm1* Ba/F3 cells were treated with MG132 (10 µM) or chloroquine (100 µM) for indicated hours, in the presence of CHX (100 µg/ml). Pde3b stability was determined by IB. Ubiquitin and Lc3b were probed to confirm the effectiveness of MG132 and CQ treatment. The Pde3b/Gapdh ratio is shown below the blots. **(K)** sgControl and sg*Ostm1* Ba/F3 cells were treated with indicated concenrations of MG132 for 8 h, and Pde3b was detected by IB. **(L)** PDE3B-His was expressed in sgControl or sg*Ostm1* Ba/F3 cells. IP with PDE3B antibody was performed, followed by IB with indicated antibodies. PDE3B-His ubiquitination was detected in sgControl cells and reduced in sg*Ostm1* cells. **(M)** OSTM1-Flag and PDE3B-His were transfected into HEK293T cells, individually or together, along with His-ubiquitin. Cells were treated with MG132 (10 µM) for 8 h prior to IP with PDE3B antibody and IB. **(N)** OSTM1-Flag and PDE3B-His were expressed separately in HEK293T cells. Cell lysates were used individually or combined for in vitro ubiquitylation assays. OSTM1-Flag promoted both its auto-ubiquitylation and PDE3B ubiquitylation. **(O)** Spearman correlation analysis of OSTM1 and PDE3B protein abundance across BCL cell lines from different subtypes using CCLE proteomic data. **(P)** OSTM1-Flag was stably expressed in indicated BCL cell lines. Flag pull-down assays were performed, and interacting proteins were analyzed by IB. **(Q and R)** OSTM1 was silenced in SU-DHL-5 (**Q**) and ARH-77 (**R**) cell lines. Two clones per cell line were probed for PDE3B by IB. **(S)** sgControl and sg*OSTM1* ARH-77 cells were treated with CHX (100 µg/ml). PDE3B was probed by IB, and its half-life was calculated based on ImageJ densitometric quantification.

The interaction between ectopically expressed OSTM1 and endogenous Pde3b was first confirmed by avidin pull-down in Ba/F3 cells expressing Flag-BirA-OSTM1 (Suppl. Fig. S4C). The OSTM1-PDE3B interaction was further validated by reciprocal co-immunoprecipitation (co-IP) in HEK293T cells co-expressing OSTM1-Flag and PDE3B-His (Fig. 4D and 4E), and at the endogenous level (Suppl. Fig. S4D). We noticed that ectopically expressed OSTM1 in HEK293T cells appeared largely glycosylated (∼60 kDa), as previously reported ^25^ and that treatment with the proteasome inhibitor MG132 increased the abundance of the non-glycosylated OSTM1 (∼37 kDa) species, which displayed enhanced interaction with PDE3B (Fig. 4D), hinting that the OSTM1-PDE3B interaction and function may be regulated by OSTM1 glycosylation status.

Next, we examined whether Ostm1 regulates Pde3b protein stability. Indeed, Pde3b protein, but not its mRNA, was increased in sg*Ostm1* Ba/F3 cells (Fig. 4F and 4G). Reconstitution of OSTM1 in sg*Ostm1* Ba/F3 cells reduced Pde3b protein levels (Fig. 4H). Cycloheximide (a translation inhibitor) chase assay showed that the half-life of Pde3b was markedly prolonged in Ostm1-silenced cells (Fig. 4I). The proteasome inhibitor MG132, but not the lysosomal inhibitor chloroquine, stabilized Pde3b (Fig. 4J and 4K). Consistent with OSTM1 functioning as an E3 ligase, its silencing decreased the ubiquitylation of PDE3B (Fig. 4L). Conversely, MG132 treatment increased K48-linked ubiquitylation of PDE3B and enhanced its interaction with the non-glycosylated form of OSTM1 (Fig. 4M). An in vitro ubiquitylation assay further demonstrated that OSTM1 directly ubiquitylates PDE3B (Fig. 4N). Analysis of Cancer Cell Line Encyclopedia (CCLE) showed an inverse correlation between OSTM1 and PDE3B protein abundance (Fig. 4O). Ectopically expressed OSTM1 and PDE3B also interacted with each other in human BCL lines (Fig. 4P). Silencing OSTM1 in the GCB-type DLBCL line SU-DHL-5 and the multiple myeloma line ARH-77 resulted in increased PDE3B protein levels (Fig. 4Q and 4R) and a prolonged PDE3B half-life (Fig. 4S). Together, these results demonstrate that OSTM1 directly interacts with PDE3B, promoting its K48-linked ubiquitylation and subsequent proteasomal degradation.

### Non-glycosylated OSTM1 preferentially interacts with PDE3B

To delineate the structural basis of the OSTM1-PDE3B interaction, we generated a series of truncation mutants of each protein, removing known or predicted functional domains. For OSTM1, we deleted the N-terminal signal peptide (SP), the luminal region containing the RING-like domain, the transmembrane (TM) domain, or the C-terminal cytoplasmic domain (CTD) ^29^. For PDE3B, we generated deletions corresponding to amino acids 1-250 and 251-650, which encompass the two N-terminal hydrophobic repeat domains, NHR1 and NHR2, respectively (Fig. 5A) ^30, 31^. Among the OSTM1 mutants, Δ306-334 that lacks the CTD, showed the most pronounced reduction in PDE3B interaction, as determined by reciprocal co-IP (Fig. 5B).

**Figure 5.**
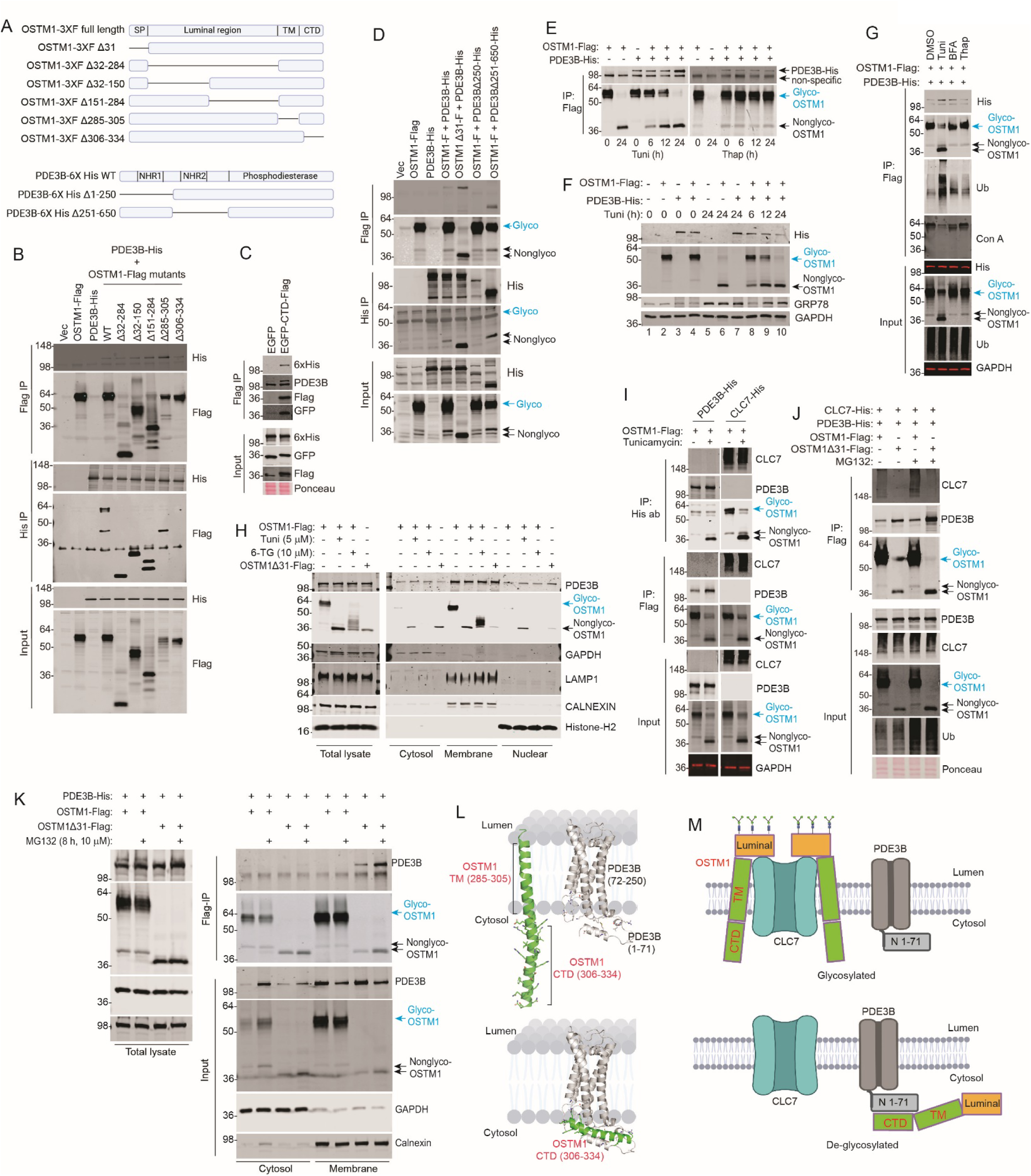
Nonglycosylated cytosolic OSTM1 interacts with PDE3B. **(A)** Schematic showing the strategy used to generate OSTM1 and PDE3B truncation mutants. OSTM1: SP (signal peptide); TM (transmembrane motif); CTD (C-terminal domain). PDE3B: NHR (N-terminal hydrophobic domain). All co-transfection assays were performed in HEK293T cells. **(B)** Cells were transfected with the indicated plasmids. IP with Flag and His antibodies was performed. The OSTM1 Δ306-334 CTD did not interact with PDE3B. **(C)** EGFP or EGFP-OSTM1-306-334-Flag was co-expressed with PDE3B-His. Flag IP shows OSTM1 CTD interacted with PDE3B. **(D)** Indicated plasmids were transfected. Reciprocal co-IP was performed using Flag and 6xHis antibodies. The N-terminal 1-250 truncation of PDE3B (Δ250) did not interact with OSTM1. **(E)** Cells co-expressing PDE3B-His and OSTM1-Flag were treated with tunicamycin (5 µg/ml) or thapsigargin (1 µM) for the indicated hours. Flag pull-down demonstrates that tunicamycin inhibited OSTM1 glycosylation and enhanced its interaction with PDE3B. **(F)** Cells co-expressing PDE3B-His and OSTM1-Flag were treated with tunicamycin (5 µg/ml) for the indicated times. Whole-cell lysates were probed for the His tag. GRP78 served as an indicator of tunicamycin-induced ER stress. **(G)** Cells co-expressing PDE3B-His and OSTM1-Flag were treated with tunicamycin (5 µg/ml), brefeldin A (10 µM), and thapsigargin (1 µM) for 24 h, followed by Flag IP. Tunicamycin increased PDE3B interaction with OSTM1. IB for ubiquitin and Con A shows increased ubiquitylation and decreased glycosylation of OSTM1 upon tunicamycin treatment. **(H)** Cells overexpressing OSTM1-Flag were treated with DMSO, tunicamycin (5 µg/ml), or 6-TG (10 µM), and cells overexpressing OSTM1Δ31-Flag were subjected to subcellular fractionation. Total cell lysates and the fractions were probed for the indicated proteins. **(I)** OSTM1-Flag was co-transfected with PDE3B-His or CLC7-His, followed by treatment with tunicamycin (5 µg/ml, 24 h). IP was performed using Flag or His antibodies. PDE3B preferentially interacted with non-glycosylated OSTM1, whereas CLC7 interacted with both glycosylated and nonglycosylated OSTM1. **(J)** Cells co-expressing CLC7-His and PDE3B-His together with either OSTM1-Flag or OSTM1Δ31-Flag were cultured in the absence or presence of MG132 (10 µM for 8 h). Flag IP was performed, followed by IB for PDE3B and CLC7. Full-length OSTM1-Flag preferentially interacted with CLC7, whereas OSTM1Δ31-Flag preferentially interacted with PDE3B. MG132 increased PDE3B and OSTM1Δ31 protein levels. **(K)** Cells expressing OSTM1-Flag or OSTM1Δ31-Flag were treated with MG132 and subjected to subcellular fractionation. Each fraction was lysed and subjected to anti-Flag IP. PDE3B was predominantly detected in membrane fractions and showed strong interaction with OSTM1Δ31. **(L)** Computational modeling of OSTM1-PDE3B interactions. Top: CHAI-predicted model of OSTM1 with its TM domain (285-305) inserted into the ER membrane and its CTD domain (306-334) oriented toward the cytosol; PDE3B residues 1-71 are cytosolic and residues 72-250 membrane-associated. Bottom: cytosolic OSTM1 allows its CTD interaction with the cytosolic N-terminal (1-71) domain of PDE3B. **(M)** Working model of OSTM1 interactions with CLC7 and PDE3B. When glycosylated and membrane-inserted, OSTM1 interacts with CLC7, positioning its RING-like domain on top of CLC7 within the luminal compartment. The transmembrane anchoring limits CTD flexibility and prevents interaction with PDE3B. Upon de-glycosylation, OSTM1 translocates to the cytosol, enabling interaction with PDE3B via the OSTM1 CTD and the cytosolic N-terminal 1-71 region of PDE3B.

Consistent with this, the OSTM1 CTD peptide (aa. 306-334), when fused to eGFP, was sufficient to directly interact with PDE3B (Fig. 5C). Because we noticed that PDE3B preferentially interacted with the non-glycosylated form of OSTM1 (Fig. 4D), we examined the glycosylation status of the OSTM1 mutants. Several luminal-region deletions, including Δ32-150, Δ151-284, and Δ285-305, remained heavily glycosylated, and in each case, PDE3B preferentially interacted with the non-glycosylated species (Fig. 5B). Importantly, OSTM1Δ31, which lacks the N-terminal 31 residues corresponding to the signal peptide^29^, was not glycosylated and exhibited strong interaction with PDE3B (Fig. 5D). Conversely, the PDE3BΔ1-250 mutant, which lacks the first NHR1 domain, showed markedly reduced interaction with OSTM1 (Fig. 5D).

To further assess whether PDE3B preferentially interacts with non-glycosylated OSTM1, we transfected HEK293T cells with Flag-tagged OSTM1 and His-tagged PDE3B, followed by treatment with tunicamycin, a pharmacological inhibitor of the first step of N-linked glycosylation. Tunicamycin treatment effectively shifted OSTM1 from its ∼60 kDa glycosylated form to the ∼37 kDa non-glycosylated species (Fig. 5E). This effect was not due to ER stress, as treatment with the ER stressor thapsigargin, an inhibitor of the sarco/ER calcium ATPases (SERCA, the ER Ca^2+^ pump), and brefeldin A (BFA), an inhibitor of the ER-Golgi transport, did not alter the glycosylation status of OSTM1 (Fig. 5E and Suppl. Fig. S5A). While tunicamycin markedly increased the co-IP between OSTM1 and PDE3B, thapsigargin failed to do so (Fig. 5E). The preferential interaction between non-glycosylated OSTM1 and PDE3B was further confirmed by in vitro binding assay using OSTM1-expressing cell lysates treated with recombinant peptide-N-glycosidase F (PNGase F), which removes N-linked oligosaccharides. PNGase F treatment resulted in OSTM1 deglycosylation and a concomitant increase in its interaction with PDE3B (Suppl. Fig. S5B). Importantly, tunicamycin treatment, which converts OSTM1 to its non-glycosylated form, also led to a marked reduction in PDE3B protein levels (Fig. 5F), suggesting that non-glycosylated OSTM1 has more potent E3 ligase activity. Indeed, tunicamycin, but not brefeldin A or thapsigargin, led to de-glycosylation of OSTM1 as indicated by reduced concanavalin A (Con A) binding, which was accompanied by elevated OSTM1 auto-ubiquitylation, a feature of active E3 ligases (Fig. 5G).

OSTM1 localizes to intracellular membranes, including the ER, trans-Golgi network, and late endosome/lysosomes, and is highly glycosylated in various tissues, including osteoclasts ^25^. The N-terminal 1-31 domain has been proposed to be a signal peptide that directs OSTM1 to intracellular membranes ^29^. Indeed, subcellular fractionation revealed that full-length wild-type OSTM1 was highly glycosylated and predominantly localized to the membrane fraction (Fig. 5H). Tunicamycin treatment profoundly inhibited OSTM1 glycosylation, resulting in a substantial proportion of non-glycosylated full-length OSTM1 redistributing to the cytosolic and nuclear fractions (Fig. 5H). In contrast, partial inhibition of OSTM1 glycosylation by the promiscuous inhibitor 6-thioguanine (6-TG) did not alter OSTM1 membrane localization (Fig. 5H).

Interestingly, OSTM1Δ31, which lacks the signal peptide, exhibited a subcellular distribution pattern similar to that of tunicamycin-treated full-length OSTM1, but was even more predominantly localized to the cytosolic fraction (Fig. 5H). The cytosolic localization of tunicamycin-treated full-length OSTM1 and the OSTM1Δ31 mutant was also observed in multiple human BCL lines (Suppl. Fig. S5C). These data indicate that glycosylation of OSTM1 critically regulates its subcellular distribution. Consequently, tunicamycin-treated full-length OSTM1 and the OSTM1Δ31 mutant are useful tools for dissecting the function of non-glycosylated and non-membrane-integrated OSTM1. Using the OSTM1Δ31 mutant, we further confirmed that the OSTM1 306-334 CTD domain is critical for its interaction with PDE3B (Suppl. Fig. S5D).

The glycosylated form of OSTM1 is believed to localize to the lysosomal membrane, where it interacts with the voltage-gated Cl^-^/H^+^ antiporter CLC7 to regulate lysosomal homeostasis in osteoclasts ^25, 32, 33^, and no E3 ligase activity was observed in this context ^29^. Since nonglycosylated OSTM1 showed a higher affinity for PDE3B (Fig. 4D, 5B, 5D-5G, Suppl. Fig. S5B), we hypothesized that OSTM1 functions are regulated by its glycosylation: the glycosylated, membrane-integrated form interacts with CLC7 to regulate ion transport and lysosomal homeostasis, whereas the nonglycosylated, cytosolic form interacts with PDE3B to promote its proteasomal degradation. Indeed, in HEK293T cells co-expressing OSTM1 together with PDE3B or CLC7, glycosylated OSTM1 preferentially interacted with CLC7, but not PDE3B (Fig. 5I, compare lanes 1 and 3). Tunicamycin treatment inhibited OSTM1 glycosylation and drastically enhanced the OSTM1-PDE3B interaction, while having little effect on the OSTM1-CLC7 interaction (Fig. 5I, lanes 2 and 4). In cells co-transfected with wild-type OSTM1 or OSTM1Δ31 together with both PDE3B and CLC7, OSTM1 was heavily glycosylated and preferentially interacted with CLC7 (Fig. 5J), whereas OSTM1Δ31 was less glycosylated and displayed a stronger interaction with PDE3B, which was further enhanced by MG132 treatment (Fig. 5J). Therefore, the OSTM1 function is critically regulated by its glycosylation and subcellular localization. While the glycosylated, membrane-integrated form interacts with CLC7, nonglycosylated OSTM1 can reside in the cytosol and interact with PDE3B to promote its ubiquitination and degradation. As PDE3B itself is also a membrane-integral protein ^30, 34^, the interaction between nonglycosylated OSTM1 and PDE3B was mainly detected in the membrane fractions (Fig. 5K).

Based on our above data and previously reported structural and subcellular localization information on the OSTM1-CLC7 complex ^32, 33^ and on PDE3B ^35, 36^, we performed computational modeling using CHAI (https://www.biorxiv.org/content/10.1101/2024.10.10.615955v2). These analyses showed that when OSTM1 is inserted in the ER membrane, its CTD domain (306-334) is unable to interact with the cytosolic (1-71) domain of PDE3B. In contrast, when OSTM1 is cytosolic, its CTD domain is predicted to engage PDE3B (Fig. 5L). We therefore propose a working model (Fig. 5M). When OSTM1 is glycosylated, it becomes membrane-tethered and interacts with CLC7, with its luminal RING-like domain positioned over the CLC7 complex in the ER lumen ^32, 33^. On the other hand, PDE3B is an integral membrane protein ^30, 34^ whose N-terminal 1-71 residues are exposed on the cytosolic side ^35, 36^. In the glycosylated state, the transmembrane (TM) domain and the CTD of OSTM1 form a rigid helical structure that sterically prevents the CTD from engaging membrane-integrated PDE3B. In contrast, upon de-glycosylation, OSTM1 translocates to the cytosol, enabling its CTD to interact with the cytosolic N-terminal 1-71 region of PDE3B (Fig. 5M).

### Nonglycosylated OSTM1 is an E3 ligase that promotes PDE3B degradation and suppresses cell growth

Given the differential glycosylation status of OSTM1 in normal human PBMCs versus human BCL lines (Suppl. Fig. S2C and S2D), we further explored the biological significance of OSTM1 glycosylation in B-cell malignancies. We first ectopically expressed full-length OSTM1 and OSTM1Δ31 in the MCL cell line Z-138 and the MM cell lines OPM2 and RPMI 8226. As observed in Ba/F3 and HEK293T cells, full-length OSTM1 was predominantly glycosylated, whereas OSTM1Δ31 was nonglycosylated (Fig. 6A). Importantly, OSTM1Δ31, but not the full-length OSTM1, reduced PDE3B protein levels and suppressed cell growth (Fig. 6A). The OSTM1Δ31-mediated downregulation of PDE3B was reversed by MG132 treatment (Fig. 6B). Consistent with our findings in HEK293T cells (Fig. 4D), MG132 also stabilized the nonglycosylated species of full-length OSTM1 in these lines (Fig. 6B), supporting the hypothesis that nonglycosylated OSTM1 possesses higher E3 ligase activity, promoting both PDE3B degradation and auto-ubiquitylation/degradation of OSTM1 itself. Similarly, full-length OSTM1 was highly glycosylated in the DLBCL cell lines OCI-LY-4, OCI-LY-7, and OCI-LY-18, and was efficiently de-glycosylated by tunicamycin treatment (Fig. 6C). While full-length OSTM1 alone did not suppress cell growth, tunicamycin treatment, which converts OSTM1 to its nonglycosylated form, profoundly inhibited the growth of cells expressing OSTM1 (Fig. 6C).

**Figure 6.**
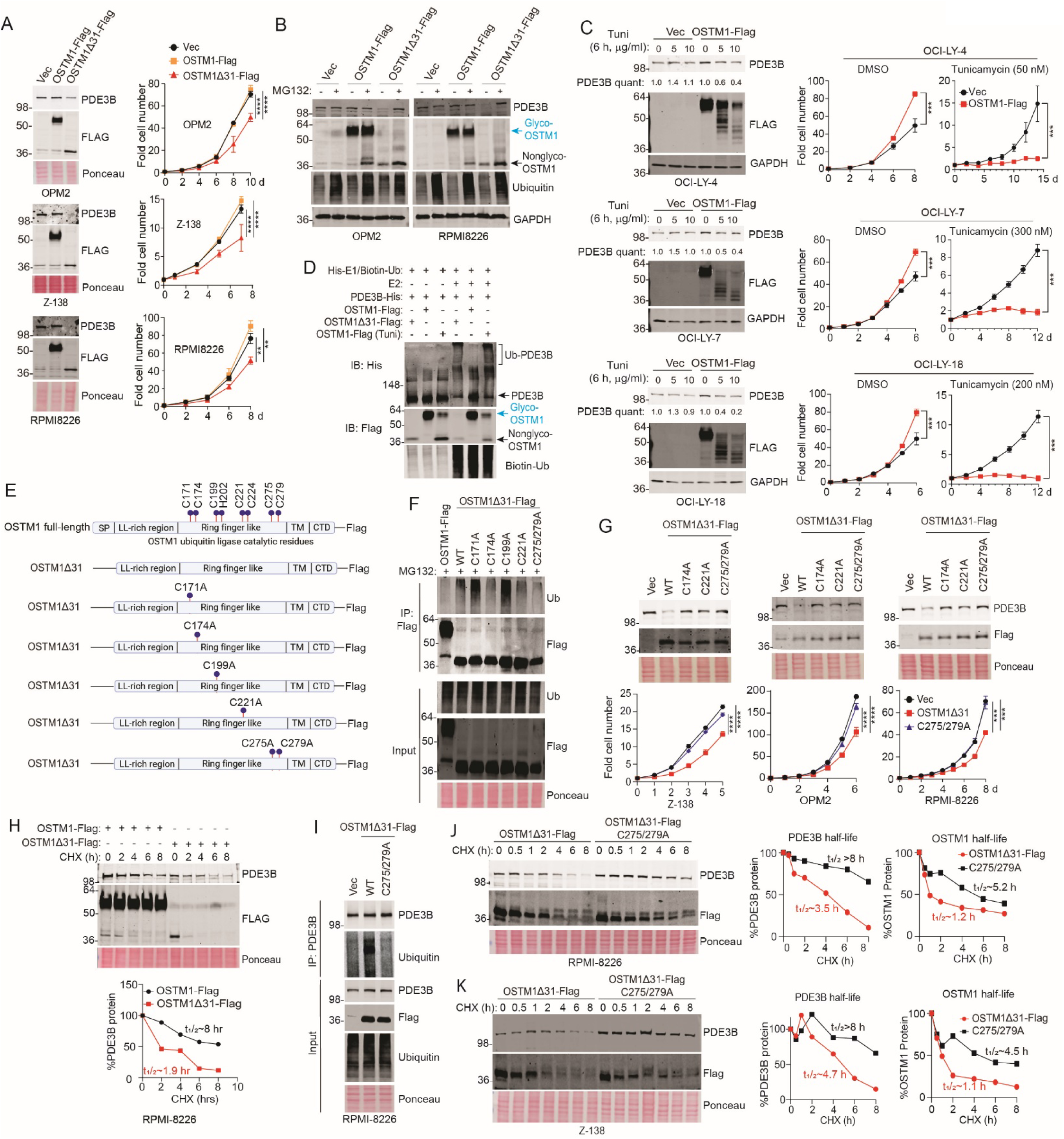
Nonglycosylated cytosolic OSTM1 is an E3 ligase that promotes PDE3B degradation and suppresses cell growth. **(A)** Wild-type full-length OSTM1-Flag or OSTM1Δ31-Flag were stably expressed in BCL lines. OSTM1Δ31, but not the full-length OSTM1, reduced PDE3B protein levels and cell growth. **(B)** BCL lines stably expressing OSTM1-Flag or OSTM1Δ31-Flag were treated with MG132 (10 µM for 8 h). OSTM1Δ31 expression led to decreased PDE3B levels, and was itself stabilized by MG132. **(C)** BCL lines stably expressing OSTM1-Flag were treated with tunicamycin. IB was performed and PDE3B level was quantified using ImageJ. Cell growth was determined by counting the number of cells. Tunicamycin led to de-glycosylation of OSTM1, decreased PDE3B stability, and resulted in more profound growth inhibition in OSTM1-expressing cells. **(D)** Cell lysates were prepared from HEK293T cells overexpressing PDE3B-His, overexpressing OSTM1-Flag treated with DMSO or tunicamycin, or overexpressing OSTM1Δ31-Flag. These lysates were used to perform in vitro ubiquitylation assays with the indicated combinations. OSTM1Δ31 and tunicamycin-treated OSTM1 lysates promoted PDE3B ubiquitylation. **(E)** Schematic showing the conserved zinc-coordinating Cys and His residues in the RING-like domain of OSTM1. The indicated Cys-to-Ala substitution mutants were generated in the OSTM1Δ31-Flag construct. **(F)** OSTM1-Flag, OSTM1Δ31-Flag and its C/A mutants were expressed in HEK293T cells. Flag IP was performed, followed by IB for ubiquitin to assess OSTM1 autoubiquitylation. OSTM1Δ31 C174A, C221A, and C275/279A mutants had markedly decreased autoubiquitylation. **(G)** OSTM1Δ31-Flag and its C/A mutants were expressed in human BCL lines. Endogenous PDE3B was detected by IB, and cell proliferation was measured. **(H)** CHX chase assay was performed in RPMI-8226 cells stably expressing OSTM1-Flag and OSTM1Δ31-Flag. PDE3B degradation kinetics was measured. **(I)** OSTM1Δ31-Flag or its C275/279A mutant was expressed in RPMI8226 cell line. PDE3B was pulled down and probed for ubiquitylation. OSTM1Δ31 enhanced PDE3B ubiquitylation, whereas the C275/279A mutant failed to do so. **(J** and **K)** CHX chase assay was performed in RPMI-8226 **(J)** and Z-138 **(K)** BCL lines stably expressing OSTM1Δ31-Flag or its C275/279A mutant. The half-lives of PDE3B and OSTM1Δ31 were determined by ImageJ densitometric analysis. Statistical significance was calculated using multiple t-tests, one per row. ***p<0.001; ****p<0.0001.

Despite that OSTM1 contains a RING-like domain ^15^, its E3 activity and biological function have remained poorly defined. To address this, we performed an in vitro ubiquitylation assay and found that both tunicamycin-treated OSTM1-wt and OSTM1Δ31 directly ubiquitylated PDE3B, indicating that nonglycosylated OSTM1 has E3 ligase activity (Fig. 6D). We then sought to further characterize the catalytic mechanism of OSTM1. Based on sequence alignment across species and conserved zinc-coordinating residues typical of RING domains (Suppl. Fig. S6) ^37^, we focused on four Cys-Cys or Cys-His pairs as candidate catalytic residues (Fig. 6E and Suppl. Fig. S6). To prevent glycosylation-mediated inhibition of E3 ligase activity, we introduced point mutations into OSTM1Δ31, rather than OSTM1-wt, to disrupt these conserved residues.

As RING-like E3 ligases typically exhibit auto-ubiquitylation, we pulled down ectopically expressed OSTM1-wt, OSTM1Δ31, and the various Cys to Ala mutants. As expected, OSTM1-wt displayed little auto-ubiquitylation, whereas OSTM1Δ31 showed robust auto-ubiquitylation (Fig. 6F). Significantly, the OSTM1Δ31 C174A, C221A, and C275/279A mutants exhibited compromised E3 activity as demonstrated by markedly reduced auto-ubiquitylation (Fig. 6F).

Consistent with this loss of catalytic activity, these mutants failed to promote PDE3B degradation, in contrast to OSTM1Δ31 (Fig. 6G). These mutants also lost the cell growth-inhibitory effect of OSTM1Δ31 in BCL cell lines (Fig. 6G). Furthermore, in RPMI8226 cells, ectopic expression of OSTM1Δ31 more profoundly accelerated the degradation of PDE3B than OSTM1-wt (Fig. 6H). Because the OSTM1Δ31 C275/279A mutant displayed the most profound loss of E3 activity (Fig. 6F), we further characterized its effect on PDE3B. Although it retained binding to PDE3B (Fig. 6I), the C275/279A mutant exhibited a prolonged half-life for both PDE3B and the mutant OSTM1 protein itself, compared with wild-type OSTM1Δ31 in both RPMI8226 and Z-138 cells (Fig. 6J and 6K). Together, these results indicate that nonglycosylated, cytosolic OSTM1 is a bona fide ubiquitin E3 ligase that interacts with PDE3B to promote its ubiquitination and proteasomal degradation.

### PDE3B suppresses cAMP/PKA/CREB signaling to promote cell growth upon OSTM1 silencing

Our findings identified PDE3B as a target for proteasomal degradation by OSTM1. PDE3B belongs to the cyclic nucleotide phosphodiesterase superfamily, which hydrolyzes cyclic adenosine monophosphate (cAMP) and cyclic guanosine monophosphate (cGMP) to produce 5’-AMP and 5’-GMP, respectively ^38^. Cytoplasmic cAMP binds to and activates protein kinase A (PKA), which in turn phosphorylates the transcription factor cAMP response element (CRE)-binding protein (CREB) ^39–42^. Phosphorylated CREB forms a transcriptional complex with CREB-binding protein (CREBBP) and the acetyltransferase EP300, initiating a tumor-suppressive epigenetic program in hematopoietic malignancies ^43–47^. Supporting the oncogenic role of PDE3B in BCL, we found that chr. 11p, where *PDE3B* is located, is amplified in nearly 17% DLBCL patients (Suppl. Fig. S7A). *PDE3B* gene gains were also observed across multiple BCL subtypes (Suppl. Fig. S7B) and were associated with increased genomic instability, as indicated by a higher fraction of genome altered (Suppl. Fig. S7C). At the mRNA level, *PDE3B* expression was elevated across several BCL patient cohorts compared with normal B cells (Suppl. Fig. S7D). At the protein level, PDE3B was low in healthy PBMCs but frequently elevated in DLBCL clinical samples (Suppl. Fig. S7E). High PDE3B expression correlated with poorer survival in both DLBCL and MM patients (Suppl. Fig. S7F).

Consistent with a pro-tumorigenic role for PDE3B, ectopic expression of PDE3B conferred IL3-independent growth in Ba/F3 cells (Suppl. Fig. S7G). Similar to observations in DLBCL patient samples (Suppl. Fig. S7E), expression of PDE3B was elevated in many BCL cell lines (Suppl. Fig. S7H). Ectopic expression of PDE3B conferred accelerated cell growth (Suppl. Fig. S7I), whereas silencing PDE3B decreased cell proliferation (Suppl. Fig. S7J and S7K) and was accompanied by increased phospho-CREB (Suppl. Fig. S7K), consistent with relief of PDE3B-mediated suppression of PKA/CREB signaling. Together, these data support a tumor-promoting role for PDE3B in B-cell malignancies.

Our findings suggest that OSTM1 suppresses BCL growth by promoting PDE3B degradation. To explore the relationship between OSTM1 and the PDE3B/cAMP/PKA/CREB1/CREBBP pathway, we first identified transcriptional target genes co-occupied by CREB1 and CREBBP by examining the available ChIP-Atlas database (https://chip-atlas.org/) (Fig. 7A). We then selected the top 1,500 CREB1/CREBBP co-occupied genes and examined their mRNA expression patterns in the TCGA MDACC BCL cohorts, correlating this gene set with the mRNA levels of CREB1, OSTM1, and PDE3B (Fig. 7B). Expression of CREBBP/CREB1 target genes showed a strong positive correlation with CREB1 and OSTM1, but a negative correlation with PDE3B across B-cell malignancies (Fig. 7C). Consistently, the CRISPR gene-effect data from DepMap (Broad Institute) revealed opposite impacts of PDE3B versus OSTM1 silencing across multiple BCL cell lines (Fig. 7D), suggesting that these two proteins exert inverse regulation within a shared signaling pathway. Together, these data support a strong inverse functional relationship between OSTM1 and PDE3B in human BCL.

**Figure 7.**
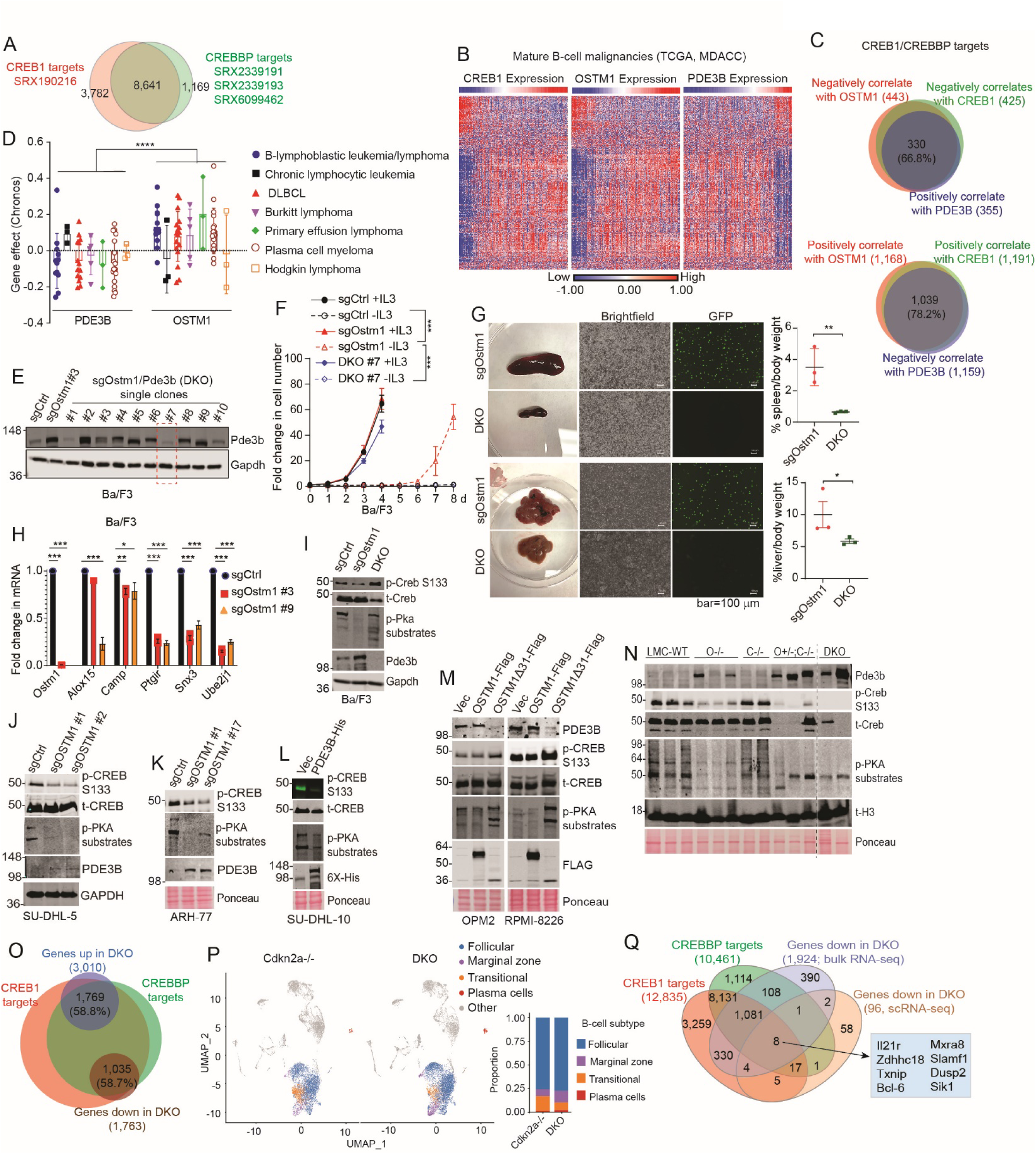
PDE3B negatively correlates with OSTM1 signaling in BCL. **(A)** Venn diagram showing overlap of B cell-specific CREB1 and CREBBP ChIP-seq targets. All targets identified in each dataset were used to determine overlap, and the top 1,500 common targets were selected for downstream analyses. **(B)** RNA expression data from the TCGA MDACC B-cell malignancies cohort. Heatmaps showing correlations between OSTM1, CREB1, or PDE3B mRNA levels and the top 1,500 CREB1/CREBBP co-occupied target genes identified in (**A**). **(C)** Venn diagrams showing overlap among genes significantly correlated with OSTM1, CREB1, or PDE3B within the top 1,500 CREB1/CREBBP targets. As OSTM1 negatively regulates PDE3B, genes positively correlated with OSTM1 and CREB1 expression and negatively correlated with PDE3B expression showed substantial overlap (78.2%). Conversely, genes negatively correlated with OSTM1 and CREB1 expression and positively correlated with PDE3B expression also overlapped significantly (66.8%). **(D)** CRISPR gene-effect scores from the CCLE database for *OSTM1* and *PDE3B* across BCL lines, showing generally negative effects upon *PDE3B* silencing and positive effects upon *OSTM1* silencing. **(E)** Pde3b was silenced by sgRNA in sg*Ostm1* clone #3 Ba/F3 cells. PDE3B protein levels were determined by IB, and sg*Ostm1/Pde3b* double knockout (DKO) Ba/F3 cells were selected for subsequent experiments. **(F)** sgControl, sg*Ostm1*, and DKO Ba/F3 cells were cutured in the presense or absence of IL3. *Pde3b* silencing reversed IL3-independence in sg*Ostm1* cells. **(G)** GFP-expressing sg*Ostm1* or DKO Ba/F3 cells were transplanted into nude mice via i.p. injection. Mice were harvested at the endpoint of the sg*Ostm1* cohort. Spleens and livers were photographed and weighed. P values were calculated using Student’s t-test. GFP-positive tumor cells were detected only in sg*Ostm1* recipients, but not in DKO recipients. **(H)** qRT-PCR in sgControl and sg*Ostm1* Ba/F3 cells showing that Ostm1 silencing reduced expression of PKA/CREB/CREBBP target genes. **(I)** sgCtrl, sg*Ostm1*, and DKO Ba/F3 cells were probed for phospho-PKA substrates and phospho-CREB (Ser133). **(J)** OSTM1 was silenced using two independent sgRNAs in SU-DHL-5 cells. Left: qPCR validation of OSTM1 knockout using on-target primers. Right: IB showing stablization of PDE3B and downregulation of cAMP/PKA signaling upon OSTM1 deletion. **(K)** OSTM1 was silenced in ARH-77 cells. IB of two clones shows increased PDE3B protein levles and decreased cAMP/PKA signaling upon OSTM1 silencing. **(L)** PDE3B-His was stably expressed in SU-DHL-10 cell line, which suppressed cAMP/PKA signaling. **(M)** OSTM1-Flag or OSTM1Δ31-Flag was stably expressed in OPM2 and RPMI-8226 cells. IB shows that OSTM1Δ31, but not the full-length OSTM1, promoted PDE3B degradation and enhanced cAMP/PKA signaling. **(N)** IB of whole-spleen lysates from indicated age-matched mice collected at the endpoints of O^+/-^;C^-/-^ or DKO cohorts. Phosphorylation levels of CREB and PKA substrates were generally reduced in the O^-/-^, O^+/-^C^-/-^, and DKO mice. **(O)** Bulk RNA-seq of purified splenic B cells from the indicated genotypes (as in **Fig. 3G**). Genes up-or down-regulated in DKO versus C^-/-^ mice were intersected with the CREB1/CREBBP ChIP-seq targets identified in **(A)**. **(P)** Spleens from two-month-old C^-/-^ and DKO mice were harvested, and 10,000 cells per mouse were analyzed by scRNA-seq. UMAP plots of B-cell subpopulations are shown by genotype. Bar graphs show the relative proportions of B-cell subsets. **(Q)** Genes down-regulated in DKO vs C^-/-^ B cells, as identified by both bulk RNA-seq of splenic B cells and scRNA-seq of follicular B cells, were intersected with CREB1/CREBBP targets. Eight genes were commonly identified across all 4 datasets.

We next directly investigated the functional connection between OSTM1 and PDE3B. Silencing Pde3b in Ostm1-silenced Ba/F3 cells reversed both IL3-independence (Fig. 7E and 7F) and in vivo transformation (Fig. 7G). Consistent with the inhibitory effect of PDE3B on the cAMP/PKA/CREB pathway, silencing Ostm1 led to reduced expression of CREB transcriptional targets (Fig. 7H) and decreased phosphorylation of CREB and PKA substrates, which were reversed by silencing Pde3b (Fig. 7I). Similarly, in human BCL cell lines, silencing OSTM1 or overexpressing PDE3B suppressed the cAMP/PKA/CREB pathway, as indicated by decreased phosphorylation of PKA substrates and CREB (Fig. 7J-7L). Importantly, OSTM1Δ31, which has stronger interaction with PDE3B and E3 activity (Fig. 5 and 6), produced a more profound decrease in PDE3B protein levels and increases in phospho-CREB and phospho-PKA substrates (Fig. 7M). Consistent with the theory that OSTM1 downregulates PDE3B to activate the cAMP/PKA/CREB pathway, Pde3b protein level was elevated in splenocytes from B cell-specific O^-/-^, O^+/-^C^-/-^, and DKO mice, which was accompanied by decreased phosphorylation of PKA substrates and CREB (Fig. 7N).

Moreover, comparison of bulk RNA-seq data from the C^-/-^ and DKO splenic B cells (Fig. 3G) revealed that 1,769 (58.8%) of the 3,010 genes upregulated and 1,035 (58.7%) of the 1,763 genes down-regulated in DKO vs C^-/-^ mice overlapped with CREB1-and CREBBP-binding targets (Fig. 7O). To further define early transcriptional changes at single-cell resolution, we performed single-cell RNA-sequencing (scRNA-seq) on spleens from young adult mice (∼2 month-old), comparing C^-/-^ and DKO genotypes. Unbiased clustering identified 8 distinct cell populations, of which approximately 51.5% were of B-cell origin (Suppl. Fig. S7L). B-cell subpopulations, including follicular, marginal zone, transitional, and plasma cells, were delineated (Fig. 7P) based on subtype-specific gene expression signatures (Suppl. Fig. S7M). At this early adult stage, 15 down-regulated genes were commonly identified by both bulk RNA-seq (Fig. 3G) and scRNA-seq and were specifically expressed in follicular B cells as determined by scRNA-seq (Fig. 7Q). Notably, 8 of these 15 DEGs were co-occupied by CREB1 and CREBBP, including *Txnip*, *Bcl-6*, *Slamf1*, *Dusp2*, *Sik1*, *Il21r*, and *Zdhhc18*, genes previously implicated in B-cell malignant transforamtion or oncogenic growth regulation (Fig. 7Q) ^48–55^.

Taken together, these data support a model in which PDE3B promotes B-cell tumorigenesis, whereas OSTM1 suppresses B-cell malignant growth by down-regulating PDE3B and activating the tumor-suppressive cAMP/PKA/CREB pathway.

## DISCUSSION

### OSTM1 is a novel TSG in BCL

We show here that in B cells, OSTM1 functions as a RING-finger ubiquitin E3 ligase that interacts with PDE3B to promote its proteasomal degradation. As PDE3B hydrolyzes cAMP and thereby limits PKA activation, OSTM1-mediated PDE3B degradation enhances cAMP-dependent PKA/CREB/CREBBP signaling, a pathway with established tumor-suppressive functions in B-cell lineages (Fig. 8). Silencing Ostm1 in the immortalized yet non-transformed murine pro-B cell line Ba/F3 induces oncogenic transformation. Importantly, although *Ostm1* ablation driven by CD19-Cre alone does not induce lymphomagenesis in naïve mice, both monoallelic and biallelic *Ostm1* deletion cooperate with Cdkn2a loss to promote malignant transformation, yielding lymphomas with histopathological features resembling multiple immature/transitional and mature B-cell malignancies.

**Figure 8.**
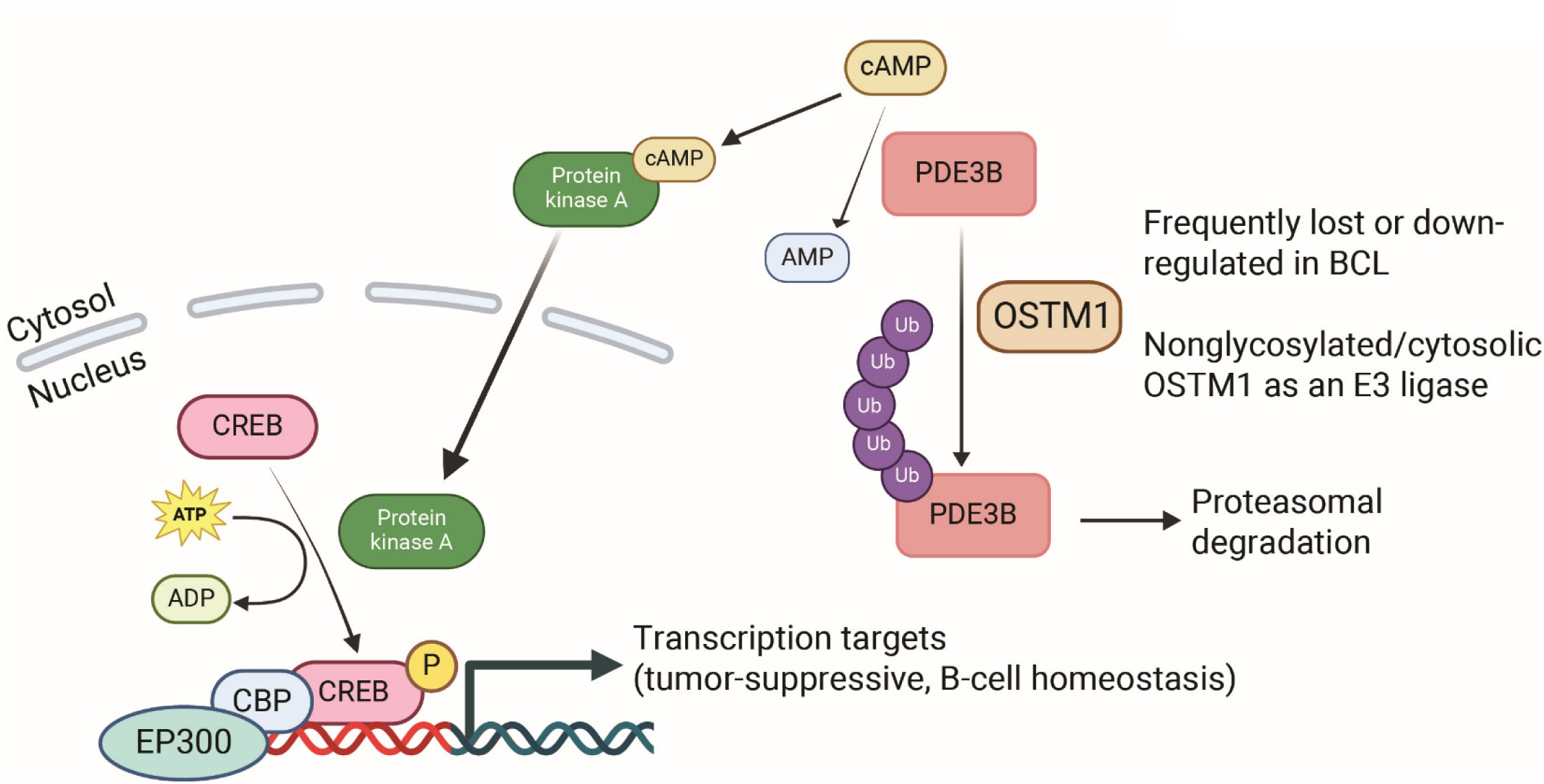
Working model: OSTM1 is a novel tumor suppressor of BCL. OSTM1 is frequently deleted in BCL due to chr. 6q loss, or its expression is otherwise downregulated, in BCL. The nonglycosylated form of OSTM1 is localized in the cytosol, where it acts as a ubiquitin E3 ligase to interact with and promote degradation of PDE3B. PDE3B hydrolyzes cAMP, thereby inhibiting the cAMP/PKA/CREB tumor-suppressive pathway. Loss of OSTM1 results in PDE3B stabilization, reduced cAMP signaling, and enhanced B-cell malignant transformation.

In humans, *OSTM1* is located on chr. 6q21. Chr. 6q arm and focal deletions are frequent events in several cancers, especially in BCLs ^18–23, 56^. Several known TSGs in B-cell lineages map to these frequently deleted regions, including PRDM1/BLIMP1 ^57, 58^, HACE1 ^59^, TNFAIP3/A20^60^, GRIK2, BACH2, CCNC, FOXO3, and EPHA7 ^61^. Previous studies have suggested that TSGs may cluster within certain chromatin regions, such that combined loss of multiple neighboring TSGs exerts additive oncogenic effects ^62^. Here we identify OSTM1 as an additional TSG in this region. Beyond genomic alterations, decreased OSTM1 expression is described in BCL patients and is associated with reduced survival (Fig. 2 and Suppl. Fig. S2). Our data further demonstrate that the E3 ligase activity and tumor-suppressor functions of OSTM1 are regulated by its glycosylation status and subcellular localization. Therefore, OSTM1’s tumor suppressor function can be negatively regulated at multiple levels, including genomic deletion, transcriptional downregulation, and post-translational modification, during malignant transformation.

### Nonglycosylated cytosolic OSTM1 functions as an E3 ligase

While OSTM1 is ubiquitously expressed, including in hematopoietic cells of the myeloid and B-and T-cell lineages, its biological and pathological roles have been described primarily in osteopetrosis and neurological disorders ^29^, mediated through its interaction with the Cl^-^/H^+^ exchanger CLC7 ^16, 25^. In this context, highly glycosylated OSTM1 interacts with CLC7 at intracellular membranes via its TM domain, and the luminal N-terminal region preceding the TM domain forms a lid-like structure above the CLC7 transporter ^32, 33^. In line with this model, pathogenic OSTM1 mutations in ARO patients are predominantly truncating or disruptive variants that abolish expression or impair proper folding of the luminal and transmembrane domains ^29^. As a result, these mutations effectively eliminate or destabilize the glycosylated, lysosomal OSTM1-CLC7 complex, indicating that glycosylated, membrane-integrated OSTM1 is required for normal osteoclast function ^32, 33^.

In addition to its interaction with CLC7 and its role in membrane trafficking, OSTM1 has been speculated to be an E3 ubiquitin ligase because it contains a RING-like domain ^15^. However, OSTM1 has not been demonstrated as a bona fide E3 ligase, and no direct E3 substrates have been reported. In fact, in the lysosomal context where OSTM1 is best characterized, the highly glycosylated OSTM1 forms a hetero-tetrameric complex with CLC7 at intracellular membranes, and its RING-like domain is localized in the luminal side of the organelle ^32, 33^. In this configuration, any E3 ligase-like activity of OSTM1 would be expected to have limited access to cytosolic ubiquitination machinery. Moreover, the acidic luminal environment and extensive N-linked glycosylation of luminal Asn residues may further constrain the catalytic activity or sterically mask the putative E3 catalytic pocket within the luminal RING-like domain.

Our results show that the PDE3B-interaction and E3 ligase activity of OSTM1 are carried out predominantly by its nonglycosylated, cytosolic form (Fig. 5L and 5M), in contrast to OSTM1’s well-recognized functions in regulating intracellular membrane trafficking and ion transport.

Because both the signal peptide and proper N-glycosylation are essential for targeting nascent proteins into the ER lumen ^63^, inhibition of N-glycosylation with tunicamycin caused a substantial fraction of OSTM1 to remain in the cytosol, where it preferentially interacts with PDE3B. Likewise, OSTM1Δ31, which lacks the signal peptide, is predominantly cytosolic and shows a strong preference for PDE3B binding (Fig. 5H and 5J). Therefore, a mechanism for OSTM1 to function as an E3 ligase in the cytosol is through its impaired glycosylation or de-glycosylation, potentially in a cell type-specific manner. Indeed, we observed a major band of nonglycosylated OSTM1 in primary human PBMCs but not in many BCL cell lines (Suppl. Fig. S2C and S2D), raising the possibility that glycosylation status may differ between normal and malignant B cells. Given that enhanced glycosylation machinery has been implicated in B-cell lymphomagenesis ^64^, increased glycosylation of OSTM1 may represent a tumor-promoting mechanism in B cells by preventing its cytosolic E3 ligase activity. It is also noteworthy that the cytosolic nonglycosylated OSTM1 is auto-ubiquitylated and degraded rapidly, which may account at least in part for the difficulty of detecting this form in previous studies. Nonetheless, our study establishes OSTM1 as a bona fide E3 ligase, a function that may have broader biological significance beyond B cells. It will be interesting to determine whether OSTM1 glycosylation status and subcellular localization vary across tissue types, especially in the other cancer types such as prostate adenocarcinoma and uveal melanoma where OSTM1 deletion frequently occurred (Fig. 2A), thereby influencing its distinct biological functions.

### OSTM1 regulates the cAMP-dependent PKA/CREB/CREBBP pathway

Our study uncovers a mechanistic connection between OSTM1 and the PDE3B/cAMP/CREB pathway. PDE3B belongs to the cyclic nucleotide phosphodiesterase superfamily and hydrolyzes cAMP to 5’-AMP ^38^. Cytoplasmic cAMP is a second messenger that regulates multiple signaling pathways by binding to its effector proteins such as PKA and EPAC ^39–42^. Upon cAMP binding, PKA is activated and phosphorylates the transcription factor CREB, which in turn drives the expression of CREB target genes. The cAMP/CREB pathway plays paradoxical roles in cancer, sometimes exerting tumor-promoting and sometimes tumor-suppressive effects, even within the same cancer type ^47^. In leukemia and lymphoma, however, cAMP signaling has largely been shown to be tumor-suppressive ^39–42, 45–47, 65^. When phosphorylated, CREB forms a complex with CREBBP and the acetyltransferase EP300, activating a tumor-suppressive epigenetic transcription program ^43, 44^.

Although transcriptional target genes of CREB1 and CREBBP have been identified by ChIP-based analyses and are available through the ChIP-Atlas database (https://chip-atlas.org/), which of these target genes are regulated by the OSTM1-PDE3B axis has remained unclear. To address this, we performed bulk RNA-seq of splenic B-cells collected at the disease endpoint of DKO mice, as well as scRNA-seq of spleens from young adult DKO and C^-/-^ mice. Cross-comparison of these four datasets enabled the identification of 15 transcriptional target genes that are co-occupied by both CREB1 and CREBBP and are consistently downregulated in follicular B cells of young adult DKO mice and in splenic B-cell lymphomas at disease endpoint. Among these, *Txnip*, *Bcl-6*, *Slamf1*, *Dusp2*, *Sik1*, *Il21r*, and *Zdhhc18* have been previously implicated in B-cell malignant transforamtion or oncogenic growth regulation ^48–55^. In this context, dysregulated expression and functions of these genes likely represent early molecular events downstream of PDE3B activation that contribute to B-cell lymphomagenesis, thereby highlighting potential pathways and targets for future mechanistic and therapeutic investigation.

In summary, our finding that OSTM1 promotes PDE3B proteasomal degradation supports a model in which OSTM1 suppresses BCL growth by relieving PDE3B-mediated inhibition of cAMP-dependent signaling and thereby enhancing PKA/CREB/CREBBP activity. These findings not only define a new layer of regulation for the cAMP/CREB/CREBBP pathway in B-cell malignancies but also expand the set of potential therapeutic targets along this axis for the treatment of BCL.

## Limitations

Clinically, OSTM1 deletions, often co-occurring with *CDKN2A* loss, are most prevalent in DLBCL and FL, particularly within the ABC-subtype of DLBCL ^66, 67^. While OSTM1 loss is also frequent in MM, *CDKN2A* loss is comparatively rare in that context. A notable limitation of our current DKO mouse model driven by *CD19-Cre* is that the resulting lymphomas predominantly originate from immature/transitional or pre-germinal center (GC) B cells. To date, we have not observed the emergence of DLBCL or FL in this model. This suggests that early oncogenic transformation in developing or pre-GC B cells may preclude the development of malignancies derived from later stages, such as GC or post-GC B cells. It would thus be interesting to further investigate the specific role of OSTM1 in mature B-cell activation contexts. This could be addressed by inducing robust GC responses through immunization of the *Ostm1*^flox/flox^*CD19*-Cre mice, or by conditionally deleting *Ostm1* in GC B cells using immunoglobulin gamma1 heavy chain (Cγ1)-Cre mice, which is active during class-switch recombination. Such approaches would enable direct testing of whether *Ostm1* loss promotes oncogenesis in GC or post-GC B cells in a manner that mirrors human DLBCL, FL, and MM pathogenesis.

## Methods

### Cell lines and culture conditions

Ba/F3 cells were kindly provided by Dr. Lina Obeid (Stony Brook University) and were cultured in RPMI-1640 (ThermoFisher, 11875119) supplemented with 10% fetal bovine serum (FBS), 1% penicillin-streptomycin (P/S) (ThermoFisher, 15140122), and 50 ng/ml interleukin-3 (IL-3) (Biolegend, 575506). Human DLBCL cell lines OCI-LY-1, OCI-LY-3, OCI-LY-4, OCI-LY-7, OCI-LY-8, OCI-LY-18, SU-DHL-6, VAL, and U2932 were cultured as previously described^68^. Human DLBCL cell lines SU-DHL-4 (CRL-2957), SU-DHL-5 (CRL-2958), and SU-DHL-10 (CRL-2963), Burkitt’s lymphoma cell lines Daudi and Ramos, as well as mantle cell lymphoma cell lines Z-138 and JeKo-1 were obtained from American Type Culture Collection (ATCC, Manassas, VA). OCI-LY-19 (ACC-528) was purchased from DSMZ. Human multiple myeloma (MM) cell lines RPMI 8226, KMS-11, and LP-1 were generously provided by Dr. Leif Bergsagel (Mayo Clinic, Scottsdale, AZ). Normal human PBMCs were obtained from the Rutgers Cancer Institute Biospecimen and Biorepository Shared Resource. All cell lines, except OCI-LY-19, were cultured in RPMI-1640 supplemented with 10% FBS, 1% P/S, 1% L-glutamine (Fisher Scientific, 25030081), 1% MEM Non-Essential Amino Acids Solution (ThermoFisher, 11140050), and 1% sodium pyruvate (ThermoFisher, 11360070). OCI-LY-19 cells were cultured in MEM-α with 90% medium, 10% FBS, and 1% P/S. All cell lines were maintained at 37°C in a humidified atmosphere with 5% CO₂. Cell lines were periodically authenticated by Short Tandem Repeat (STR) profiling at IDEXX BioResearch.

### Antibodies

Flag-M2 (Millipore Sigma, F1804), Flag (Millipore Sigma, F7425), 6X-His (Thermo Fisher Scientific, MA1-21315), p-AKT Ser-473 (Cell Signaling Technology, 4060), t-AKT (Cell Signaling Technology, 9272), GAPDH (ProteinTech, 10494-1-AP), p-S6 Ser235/236 (Cell Signaling Technology, 4858), t-S6 (Cell Signaling Technology, 2217), p-STAT5 Y694 (Cell Signaling Technology, 9359), t-STAT5 (Cell Signaling Technology, 25656), p-p44/42 MAPK (Erk1/2) Thr202/Tyr204 (Cell Signaling Technology, 4376), t-p44/42 MAPK (Erk1/2) (Cell Signaling Technology, 9102), PDE3B (SMCP3B) (used for IB) (Novus Biologicals, NBP1-43333), PDE3B (used for IP) (Abcam, ab99290), Ubiquitin (P37) (Cell Signaling Technology, 58395), LC3B (Cell Signaling Technology, 2775), K48-linkage specific polyubiquitin (D9D5) (Cell Signaling Technology, 8081), GFP (Santa Cruz, sc-8334), BiP/GRP78 (BD Biosciences, 610978), calnexin (Transduction Labs, C45520-050), CLC7 (Thermo Fisher, A305-381A-T), p-CREB Ser133 (87G3) (Cell Signaling Technology, 9198), t-CREB (48H2) (Cell Signaling Technology, 9197), p-(Ser/Thr) PKA substrates (Cell Signaling Technology, 9621), OSTM1 (Millipore Sigma, ABN1365), goat anti-mouse IgG (H+L) cross-adsorbed secondary antibody, Alexa Fluor 680 (Thermo Fisher, A-21057), goat anti-rabbit IgG (H+L) cross-adsorbed secondary antibody, Alexa Fluor™ 594 (Thermo Fisher, A11012), goat anti-rabbit IgG (H+L) cross-adsorbed secondary antibody, Alexa Fluor 488 (Thermo Fisher, A11008), goat anti-rabbit secondary antibody (LI-COR, 926-32211), goat anti-rat secondary antibody for 680 channel (Thermo Fisher, A-21096)

### Reagents, chemicals, and kits

6-thioguanine (6-TG) (MedchemExpress, HY-13765), biotin (Millipore Sigma, PHR1233), brefeldin A (Thermo Fisher, 00-4506-51), chloroquine (Selleckchem, S6999), concanavilin A (Vector Labs, B1005-5), cycloheximide (Millipore Sigma, C4859), L-glutamine (Fisher Scientific, 25030081), IRDye 800CW streptavidin (LI-COR Bio, 926-32230), IL-3 (mouse recombinant, carrier-free. BioLegend, 575504), leupeptin hemisulfate (MedchemExpress, HY-18234A), MG132 (Selleckchem, S2619), MEM non-essential amino acids solution (Thermo Fisher, 11360070), NGI-1 (MedchemExpress, HY-117383), Protein G Dynabeads™ (Thermo Fisher, 10004D), sodium pyruvate (Thermo Fisher, 11360070), thapsigargin (Millipore Sigma, T9033), TRIzol™ Reagent (Thermo Fisher, 15596018), tunicamycin (Tocris Bioscience, 3516), polyethylenimine (PolySCIENCES, 23966-1), ubiquitylation kit (Enzo Life Sciences, BML-UW9920-0001).

### Constructs

For stable and transient overexpression, OSTM1-3X-Flag, PDE3B-6X-His and CLC7-6X-His were cloned into pLV-EF1a-IRES-Neo (Plasmid #85139 in Addgene). For mouse and human OSTM1 and PDE3B genes CRISPR/Cas9 based knockout, pLenti-CRISPR-V2-puro plasmids with cloned sgRNAs were ordered from Genscript (2 guides/gene/organism). hOSTM1 (guide 1, TGAACATCACTCGAAAACTA; guide 2, CTGCTTTGAACATAACCTTC), hPDE3B (guide 1, TACTGACTATCCCGAAGCAA; guide 2, TACTGACTATCCCGAAGCAA), mOstm1 (guide 1, TCGATGCACAAGTGCGTCTC; guide 2, GCTATTGCAGAATACCTCCG) mPde3b (guide 1, GCTAAATGAAGCTCGCAATA; guide 2, GGGATCGCAGCAGTGGTAAG). His-ubiquitin, HA-ubiquitin, and Flag-BirA-OSTM1 were cloned into LPC vector. OSTM1-CTD (306-334)-Flag peptide was cloned into pEGFP-C3. All plasmids were purified using Plasmid Maxi Kit (Qiagen, 12163) and confirmed by Sanger sequencing at GeneWiz.

### Lentivirus generation and infection

For lentivirus generation to stably overexpress or knockout our genes of interest, PSPAX2 (packaging plasmid), pVSVG (helper plasmid) and target plasmids (pLV-EF1a-IRES-Neo/Puro or pLenti-CRISPR-V2-puro) were transfected into HEK293T cells at 1 (PSPAX2): 3 (pVSVG): 4 (target plasmid) ratio using polyethylenimine (PEI) transfection method at 1 (DNA): 3 (PEI) ratio. After overnight incubation, media were changed to fresh media and 48 h later supernatant was collected to infect target cells at 1×10^6^ cells/well density with 5 μg/ml polybrene in 12-well plate. Plate was centrifuged at 1,000 rpm for 2 h at room temperature, then incubated overnight in CO_2_ incubator. Culture media were then changed back to fresh media for respective cell lines, followed by antibiotic selection or cell growth measurement.

### Cell proliferation assay

Cells were seeded at different densities (Ba/F3 1,000, SU-DHL-5 1,000, OPM2 2,000, Z-138 2,000, RPMI-8226 1,000, ARH-77 1,000, OCI-LY-4 5,000, OCI-LY-7 3,000, OCI-LY-18 2,000, U2932 1,000 and SU-DHL-10 1,000 cells per well) in 96-well plates. Cell number was recorded each subsequent day using Celigo High Throughput Micro-Well Image Cytometer (NEXCELOM Bioscience, Art nr: 200-BFFL-5C-AUTO) until maximum confluence was reached. Fold change in cell number was calculated by normalizing the cell count of each day to that of day 0.

### Drug treatments for co-IP and IB

Cells (both adherent and suspension) were treated with MG132 (10 µM or dose response), chloroquine (100 µM), tunicamycin (5 µg/mL), 6-thioguanine (10 µM), thapsigargin (1 µM) or NGI-1 (10 µM) for the indicated time points. Following treatment, cells were collected for co-immunoprecipitation (co-IP) and/or immunoblotting (IB) analysis or subjected to subcellular fractionation followed by IB.

### Cell growth and proliferation assay

Cell growth and proliferation were assessed by daily cell counting. Equal number of cells between tests and controls were seeded in 96-well plates. Cell numbers were quantified daily using a Celigo cell counter. Cell counts were normalized to day 0 values, and proliferation curves were plotted in GraphPad Prism 8.

Cell viability was additionally assessed using trypan blue exclusion. The percentage of viable and non-viable cells was determined daily, normalized to that of day 0.

### Drug sensitivity assay

For drug sensitivity assays, an MTS assay was performed. Cells were seeded in 96-well plates and treated with increasing concentrations of each inhibitor for 72 h, with DMSO-treated cells serving as controls. At the endpoint, MTS reagent (Boc Sciences, B2699-049332) and phenazine methosulfate (PMS; Millipore Sigma, P9625) were mixed and added to each well for 3-4 h or until color development was evident. Absorbance was measured at 490 nm using a Molecular Devices M5 plate reader. Percent viability was calculated relative to untreated controls, and IC₅₀ values were determined by non-linear regression (four-parameter logistic curve) using GraphPad Prism 8.

### De-glycosylation co-IP in cultured cells

For in-cell de-glycosylation and subsequent co-IP for determination of the effect of OSTM1 glycosylation on OSTM1-PDE3B interaction, OSTM1-Flag and PDE3B-His were co-transfected in HEK293T cells. Cells were treated with 5 µg/ml tunicamycin or 10 µM NGI-1 for 24 h. Cells were lysed, and 500 µg of protein lysate was co-immunoprecipitated with Flag-M2 magnetic beads and eluates were utilized for IB to evaluate the effect OSTM1 glycosylation on its interaction with PDE3B or CLC7.

### In-vitro de-glycosylation co-IP

For investigating the role of OSTM1 glycosylation on OSTM1-PDE3B interaction, OSTM1-Flag and PDE3B-His were separately overexpressed in HEK293T cells. OSTM1-Flag expressing cell lysates underwent in-vitro PNGase-F treatment (NEB, P0704) under non-denaturing conditions following manufacturer’s protocol. Briefly, lysates were incubated with PNGase-F for 4 h at 37^°^C. PNGase-F treated OSTM1-Flag expressing cell lysates were co-incubated with PDE3B-His expressing cell lysates along with Flag-M2 or 6x-His magnetic beads to purify complexes, followed by IB.

### Ubiquitylation assay in cultured cells

#### Non-denaturing IP-ubiquitylation detection

Stable OSTM1-Flag or PDE3B-His overexpressing (Ba/F3 or BCL cell lines) or transiently transfected (using PEI method) HEK293T cells were lysed in IP lysis buffer, and 500 µg protein lysate was incubated with Protein G beads and respective antibodies at 4^°^C overnight. For endogenous PDE3B pull down, PDE3B-ab was used. For cells overexpressing PDE3B-His, 6x-His tag antibody was used. Beads were separated by magnetic separation and washed 2x with wash buffer 1, 2x with wash buffer 2 and once with IP lysis buffer. Bound PDE3B was eluted with 1x Laemmli buffer, followed by IB with ubiquitin antibody.

#### High-denaturing IP-ubiquitylation detection

1×10^6^ HEK293T cells were plated into 6-cm petri dish. After overnight attachment, cells were transfected with 2 µg HA-Ub, 2 µg PDE3B-His, and 2 µg OSTM1-Flag constructs using PEI transfection. Cells were harvested 48 h after transfection. One tenth of cells were lysed in RIPA buffer and the remaining cells were dissolved in Buffer A (6 M guanidine-HCl, 0.1 M Na_2_HPO_4_/NaH_2_PO_4_, 10 mM imidazole, pH8.0) and sonicated. Each sample was added with 50 µl PureCube Ni-NTA MagBeads (Cube Biotech, 31205) equilibrated with Buffer A and incubated for 3 h at room temperature on rotation. The agarose beads were precipitated by centrifugation and washed once with Buffer A, twice with Buffer B (10 mM Tris-Cl, pH 8.0, 8 M urea, 0.1 M NaH_2_PO_4_) and three times with 1:4 diluted Buffer B supplied with 25 mM imidazole. The precipitates were resuspended in 100 µl 2Х Laemmli buffer (4% SDS, 20% glycerol, 10% β-mercaptoethanol, 0.004% bromophenol blue, 0.125 M Tris HCl pH 6.8) with 200 mM imidazole, boiled at 95℃ for 10 min and subjected to IB.

#### In-vitro ubiquitylation

For in-vitro ubiquitylation of PDE3B in the presence of OSTM1, in-vitro ubiquitylation kit (Enzo Biochem., BML-UW9920-0001) was used following manufacturer guidelines. In brief, OSTM1-Flag, OSTM1Δ31-Flag, or tunicamycin treated OSTM1-Flag expressing cell lysates were co-incubated with PDE3B-His, E1, E2, Mg-ATP, and streptavidin-tagged ubiquitin for 2 h. Resultant reactions were used for IB.

#### Cancer Cell Line Encyclopedia (CCLE) data analysis

CRISPR-Cas9 gene dependency (Chronos Gene Effect) and protein abundance data for OSTM1 and PDE3B were obtained from the CCLE database (DepMap Public 23Q4+ release; https://depmap.org/portal/). Gene effect scores were used to assess dependency across BCL cell lines and visualized as bar graphs in Graphpad Prism 8.

Protein abundance data, expressed as relative abundance normalized to the bridge sample, were available for a subset BCL cell lines. Pearson correlation between protein abundances of OSTM1 and PDE3B was calculated in Microsoft Excel, and correlations were visualized as scatter plots in GraphPad Prism 8.

### Publicly available patient data analysis

#### Copy number analysis

For *OSTM1*, *CDKN2A,* and *PDE3B* copy number analysis across B-cell malignancies, two datasets were available in cBioPortal: DLBCL (Lymphoid Neoplasm Diffuse Large B-cell Lymphoma (TCGA, Firehose Legacy) with an n=48 and Mature B-cell malignancies (MD Anderson Cancer Center) with 752 patients spread across different B-cell lymphoma subtypes such as DLBCL, MCL, FL, BL, MZL, HGBL and CLL). The copy number data was obtained from these datasets to determine the frequency of alterations. Co-alteration analysis was done by making oncoplots using Oncoprinter (https://www.cbioportal.org/oncoprinter).

Furthermore, copy number analysis was also performed for several other publicly available datasets ^20^ (n=304), multiple myeloma ^23^ (n=725), B-cell chronic lymphocytic leukemia ^19^ (n=97), follicular lymphoma ^22, 61^, plasmablastic lymphoma ^21^, and Burkitt’s lymphoma ^18^.

#### Analysis of publicly available GEO datasets

Gene expression datasets relevant to BCL were retrieved from the NCBI Gene Expression Omnibus (GEO). For each dataset, the series matrix file and the corresponding platform annotation file were downloaded. Expression data were aligned to the annotated gene symbols and combined for downstream analyses.

For *OSTM1*, tumor versus normal expression was analyzed using GSE32018, GSE56315, and GSE12195, while survival analyses were performed using GSE10846, GSE181063, GSE4475, GSE119214, and GSE2658. For *PDE3B*, tumor versus normal expression comparisons included GSE2350, GSE56315, GSE12195, and GSE50006, and survival analyses were conducted using GSE4475 and GSE2658.

All analyses were performed using Microsoft Excel. Expression differences between two groups were assessed using Student’s *t*-test, and differences among more than two groups using one-way ANOVA. For survival analysis, patients were stratified into high-and low-expression groups based on the median expression of the indicated gene, and survival plots were generated accordingly.

#### Correlation analysis of CREB1–CREBBP target genes

CREB1 and CREBBP target genes were identified from ChIP–seq datasets available in ChIP-Atlas (CREB1: SRX190216; CREBBP: SRX2339191, SRX2339193, SRX6099462). The top 1,500 genes common across these datasets were selected for further analysis. Their expression patterns were examined in the cBioPortal “Mature B-cell Malignancies (MD Anderson Cancer Center)” dataset. Pearson correlation coefficients between CREB1/CREBBP target genes and OSTM1, PDE3B, and CREB1 expression were calculated in Microsoft Excel. Statistically significant correlations (p < 0.05) were visualized using Morpheus, and overlapping gene sets (e.g., PDE3B negatively correlated vs. OSTM1 and CREB1 positively correlated genes) were identified using the Venny tool.

#### Ba/F3 cell line derived allograft mice

sgControl and sg*Ostm1* Ba/F3 cells were GFP labelled using lentiviral infection. 1×10^6^ GFP labeled cells were resuspended in 500 μl DPBS (Thermo Fisher, 14190250) and injected via intraperitoneal (i.p.) in nude mice. Mice were monitored until the development of tumor related characteristics such as abdominal enlargement and lethargy. At the endpoint, survival span was recorded, mice were euthanized, and secondary lymphoid organs spleen, liver, and lymph-node weights were recorded. Subsequently, tumorigenic organs were crushed, and single cell suspension was prepared in DPBS and analyzed under fluorescent microscope for GFP^+^ cells enrichment comparison between sgControl and sg*Ostm1* allografts.

#### Cycloheximide chase assay

2×10^6^ cells (ARH-77, Ba/F3, RPMI-8226, and Z-138 cells) were seeded in 12-well plates and treated with 100 µg/ml cycloheximide or DMSO. Cells were collected at different timepoints, washed with DPBS, lysed in 2x Laemmli buffer then subjected to IB.

#### Tandem mass tag (TMT) proteomics

Cell pellets from sgControl (triplicate) and sg*Ostm1* (triplicate) were used for protein identification and relative quantitation using multiplex TMT approach. Total protein was extracted using lysis buffer containing 8 M urea, 100 mM TEAB and 1X protease inhibitor with sonication. Protein concentration was measured using BCA assay. Fifty microgram total protein from each replicate was reduced by DTT and alkylated by IAM followed by in-solution trypsin digestion.

The resulting peptides were labeled with TMT reagents. After the TMT labeling, the peptides from all 6 samples were combined and desalted using C18 cartridge followed by high pH RPLC separation. A total of 12 fractions were collected and further analyzed by RPLC-MS/MS on an Orbitrap Fusion Lumos Mass Spectrometer using MS2 method. The MS/MS spectra were searched against UniProt mouse database using Sequest search engine on Proteome Discoverer (V2.4) platform. The quantitation comparison was calculated as follows:

Ratio (sg*Ostm1*/sgCtrl) = The sum of (each group) TMT abundance/The sum of (sgCtrl) TMT abundance. The proteins with the ratios greater than 1.5-fold and passed the T-Test (p≤0.05) are considered significantly changed and used for further analysis.

#### Bio-ID mass spectrometry

Flag-BirA-OSTM1 transiently or stably expressing cells were grown in the presence of 50 μM biotin for 24 h. Cells were harvested and lysed in RIPA buffer (50 mM Tris (pH 7.4), 150 mM NaCl, 1% Triton X-100, 1% sodium deoxycholate, 1% SDS) with sonication. 500 µg of total lysate was incubated with 100 µl avidin beads slurry at 4^°^C overnight with rotation. Avidin beads were washed twice with wash buffer 1 (2% SDS in dH_2_O), once with wash buffer 2 (20 mM Tris, pH 7.4, 500 mM NaCl, 1% Triton X-100, 1 mM EDTA, 1 mM EGTA), once with wash buffer 3 (20 mM Tris, pH 7.4, 300 mM NaCl, 1% Triton X-100, 1 mM EDTA, 1 mM EGTA) and once with wash buffer 4 (20 mM Tris, pH 7.4, 100 mM NaCl, 1% Triton X-100, 1 mM EDTA, 1 mM EGTA). Bound proteins were removed with SDS sample buffer boiling at 95^°^C for 5 min. Eluates were run on SDS-PAGE, then subjected to IB or mass-spectrometry (MS).

For MS, in-gel trypsin digestion was performed. The resulting peptides were analyzed by LC-MS/MS on Orbitrap Fusion Lumos MS instrument. The MS/MS spectra were searched against Uniprot mouse database using Sequest search engines on Proteome Discoverer (V2.4) platform. The protein false discovery rate is less than 1%. The ratio of Flag-BirA-OSTM1/vector was calculated. The proteins containing more than one unique peptide are identified with more confidence.

#### Site-directed mutagenesis (SDM)

SDM was performed using Q5-SDM kit by New England Biolabs (NEB) following manufacturer’s instructions. All mutagenesis was performed on constructs cloned in pLV-EF1a-IRES-Neo vector. NEBaseChanger was used for designing mutagenesis primers. For point mutants (substitutions), mutations were introduced in the forward primer, used in the SDM PCR reaction. For truncation mutants, forward and reverse primers were made such that the 5’ tails of both forward and reverse primers were placed adjacent to the target deletion region for running the PCR reaction.

#### RNA extraction, cDNA synthesis, qPCR

RNA of tissue/cell lines was isolated using TRIzol™ Reagent based on manufacturer’s guideline. Subsequently, reverse transcription was done using Superscript III First Strand Synthesis system (Invitrogen). qRT-PCR was performed with Power SYBR™ Green PCR Master Mix (Thermo Fisher) on StepOnePlus (applied following primers: mPde3b-F: GCTAAATGAAGCTCGCAATA mPde3b-R: GTGGAGTTGGGAAACTGGT; mOstm1 (on-site)-F: CGAACTGCGCAAATTGCCT, mOstm1 (on-site)-R: TCGATGCACAAGTGCGTCTC; hOSTM1 (mRNA expression)-F: AAGATGCAATGAACATCACTCG, hOSTM1 (mRNA expression)-R: GACTTGAGACGTTTGGGCAG; hOSTM1 gRNA1 (on-site)-F: CTGCTTTGAACATAACCTTCAGGG, hOSTM1 gRNA1 (on-site)-R: GTCACTGCAAGGGACTGAAC; hOSTM1 gRNA2 (on-site)-F: GCCAGAAGTCTCTTAATGGCAGAT, hOSTM1 gRNA2 (on-site)-R: TGTGCATTCCCCTGAAGGTT; mAlox15-F: GACACTTGGTGGCTGAGGTCTT, mAlox15-R: TCTCTGAGATCAGGTCGCTCCT; mCamp-F: CTTCAACCAGCAGTCCCTAGAC, mCamp-R: GCCACATACAGTCTCCTTCACTC; mPtgir-F: GTTTACCACCTGATTCTGCTGGC, mPtgir-R: CGTTGAAGCGGAAGGCGAGGA; mUbe2j1-F: GAAACCTGGCAGCCTTCATGGA, mUbe2j1-R: GCCTTCACAACAGAAATCTTGCG; mSnx3-F: ACAGTGACTTTGAGTGGCTTCGA, mSnx3-R: CCTCTAAAAGGAAGCTGCCGCA,

#### Genomic DNA sequencing for CRISPR knockout validation in cell lines

Genomic DNA (gDNA) was isolated from sgControl and sg*OSTM1* ARH-77 and Ba/F3 cells using QIAamp DNA Mini Kit (Qiagen, 51304). Isolated gDNA was used to setup PCR using primers designed from region flanking gRNA targeting mouse and human OSTM1 for knockout. For Ba/F3 cells, following primer set was used: F: CACGCAGAGATACTGTGTCCA, R: ACAGAGACACCGTAGAGCCT and for ARH-77 cells, following primers were used: F: ACTGTAGTGTGAAACGAAGCTG, R: ACAATCACACACCTCAATCAATGT for amplifying ∼0.5kb size amplicon. The amplicons were used for sanger sequencing using sequencing primer CATACAGTCTCCTCCCACCAAA for Ba/F3 and ATATAAATGACAACATAGAATTGTTC for ARH-77 cells.

#### Co-immunoprecipitation

Adherent (HEK293T) or suspension (BCL cell lines) cells were collected and centrifugated at 1,000 x g for 3 min. After being washed with cold PBS, cells were lysed in IP lysis buffer (20 mM Tris, pH 7.5, 100 mM NaCl, 1% Triton X-100, 1 mM EDTA, 200 mM PMSF) supplemented with protease inhibitor cocktail on ice for 30 min. The cell lysates were cleared by centrifugation at 4°C to collect supernatant. The supernatant (250-500 μg) was incubated with magnetic beads (Anti-FLAG M2) or Protein G Dynabeads™ with either 6x-His or PDE3B antibodies) overnight at 4°C on rotator. The complexes were precipitated with magnetic stand, and washed twice with IP wash buffer 1 (20 mM Tris, pH 7.5, 300 mM NaCl, 1% Triton X-100, 1 mM EDTA, 200 mM PMSF) in a rotator for 5 min, washed twice with IP wash buffer 2 (20 mM Tris, pH 7.5, 200 mM NaCl, 1% Triton X-100, 1 mM EDTA, 200 mM PMSF) for 5 min, then once with IP lysis buffer for 5 min, and resuspended the complexes with 60 μl IP lysis buffer, boiled in SDS sample buffer at 95°C for 5 min and subjected to IB.

#### Immunoblotting using cell lysate from cell lines and mouse tissues

Spleen tissues were harvested and snap frozen in liquid nitrogen. Lysates were prepared using homogenization in RIPA buffer with 1% SDS and protease inhibitors cocktail (MCE Chemicals) and quantified using BCA kit (Thermo Fisher, A55864). BSA was used for making standard curves. For cell lines, lysates were prepared in 2X Laemmli sample buffer (1610737EDU) and quantified using Pierce™ 660nm Protein Assay Reagent (Thermo Fisher, 22660) with the addition of Ionic Detergent Compatibility Reagent for Pierce™ 660nm Protein Assay Reagent (Thermo Fisher, 22663). 10-30 μg protein was separated in SDS-PAGE and transferred to nitrocellulose blotting membrane. Primary antibodies were incubated overnight at 4 °C in 5% milk in TBST, and Alexa fluor conjugated secondary antibody was incubated for 1 h at room temperature in 5% milk. Imaging the blot was done using the Odyssey imaging system (LI-COR Biosciences).

#### Mouse experiments

*Cdkn2a*^flox/flox^ mice (B6.129S1-*Cdkn2a*<tm4Cjs>/J, Strain #: 023323), and *Cd19*^+/Cre^ mice (B6.129P2(C)-*Cd19^tm1(cre)Cgn^*/J) were purchased from the Jackson Laboratory. *Ostm1*^lox/lox^ mice (B6;129-*Ostm1^tm1.1Jva^*/J) were generated in Jean Vacher laboratory ^26^. Experimental mice were produced by breeding *Ostm1*^flox/flox^, *Cdkn2a*^fflox/flox^ mice, and *Cd19*^+/Cre^ mice to produce wild-type littermate control (WT-LMC), O^-/-^, C^-/-^, O^+/-^C^-/-^, and O^-/-^C^-/-^ (DKO) mice. Age-and gender-matched both male and female mice were used. For the Ba/F3 allograft and transplantation experiment, athymic nude mice (*Foxn1*^nu^/*Foxn1*^nu^) were purchased from the Jackson Laboratory. Six to eight-week-old female nude mice were used. All mouse experiments were performed in compliance with the Institutional Animal Care and Use Committee guidelines at Rutgers University.

#### Flow cytometry

At endpoint, two femur bones were isolated and separated from attached muscular tissue. The ends were cut and the marrow was flushed in HBSS buffer. Spleen tissue was physically disrupted between frosted slides to get a single cell suspension. Splenocytes were washed with HBSS and subject to ACK (Thermo Fisher, A1049201) lysis to remove RBCs and passed through a 70 μm cell strainer (Fisher Scientific, 22-363-548). 2 x 10^6^ cells from bone marrow and splenocytes were blocked with rat serum and FcR blocking antibody 2.4G2 (BioLegend 101302) and incubated with fluorophore conjugated antibodies: Cd3e (BioLegend 100351), B220 (Biolegend 103206), CD21/CD35 (BioLegend 123434), CD23 (BioLegend 101621), IgD (BioLegend 405720), IgM (BioLegend 406509), CD38 (BioLegend 102720) CD138 (BioLegend 142534), Gr-1 (BioLegend 108456), CD93 (BioLegend 136503). Data were acquired on Cytek Aurora Spectral Flow Cytometer (Cytek Biosciences, Fremon, CA) and analyzed on the FlowJo software (TreeStar, San Carlos, CA).

#### Bulk-RNA sequencing of purified B cells and data analysis

Splenic B cells were purified from WT, *Ostm1^-/-^, Cdkn2a^-/-^* and *Ostm1^-/-^;Cdkn2a^-/^*^-^ (DKO) mice using CD43 negative selection. Briefly, spleen was harvested from mice and single cell suspension was prepared. Red blood cell lysis was done using ACK lysing buffer (Thermo Fisher, A1049201) and the cell suspension was passed through 70 µm strainer (Fisher Scientific, 22-363-548).

Splenocytes were incubated with microbeads coated with anti-CD43 antibody (Miltenyi Biotec, 130-049-801) and passed through LS columns (Miltenyi Biotec, 130-042-401). Resultant eluates containing splenic B cells were counted using trypan blue counting and subjected to RNA isolation. RNA purity was confirmed by running 1% agarose gel. RNA-sequencing was performed at the University of Texas Health San Antonio (UTHSA) Genome Sequencing Facility. RNA-seq library was prepared with 500 μg of total RNA following the Illumina TruSeq stranded mRNA sample preparation guide. After quantification, the library was subjected to cBot amplification and subsequent 50 bp single read sequencing on the Illumina HiSeq 3000 platform using 100 PE sequencing module.

The RNA-seq reads were mapped to the mouse genome reference (Mus musculus genome (UCSC mm10)) using HISAT2, Gene-level quantification was counted using feature counts, differentially expressed gene analysis was conducted using DEseq, heatmap was generated using iDEP online tool https://bioinformatics.sdstate.edu/idep/. Functional enrichment analysis was performed using the Metascape online tool https://metascape.org/gp/index.html#/main/step1 and KEGG pathway analysis using ShinyGO 0.85 https://bioinformatics.sdstate.edu/go/.

### Single-cell RNA sequencing (scRNA-seq)

#### Cell Isolation

Spleens were harvested from experimental mice and placed in cold DPBS. Tissues were immediately cut in small pieces and then gently pressed through a 100 µm mesh cell strainer using the plunger end of a syringe. Cell suspension was diluted with DPBS and centrifuged at 400 x g for 5 min at 4°C. To eliminate red blood cells, the cell pellet was resuspended in 5 mL of red cell lysing buffer (Roche Diagnostics) and incubated for 3 min at room temperature. Cell suspension was then filtered through a 30 µm mesh cell strainer, diluted with cold DPBS, and centrifuged. After additional washing with DBPS, splenocytes were resuspended in DPBS containing 0.4% (w/v) bovine serum albumin (BSA) (Millipore Sigma) and counted using a hemocytometer with cell viability assessed using Trypan Blue staining. A sample containing 1.5 x 10^6^ viable cells/mL was filtered through a 20 µm mesh cell strainer, counted again, and then used for single cell RNA-sequencing. Cell viability for all samples ranged between 80 and 92%.

#### Library construction and sequencing

Single cell RNA-sequencing was performed by the Stony Brook University/Northport VAMC Single Cell and Spatial Multiomics Facility, using the Chromium Next GEM Single Cell 5’ Reagent kit v2, according to the manufacturer’s instructions (10x Genomics, protocol CG000331). Briefly, 10,000 single cells per sample were captured in Gel Bead-In-Emulsions (GEMs) using Chromium iX (10x Genomics). After reverse transcription in a PCR cycler, the barcoded full-length cDNA was purified, amplified with PCR for 11 cycles, and further purified with SPRISelect reagent (Beckman Coulter). Fragment size distribution and quantification of the amplified cDNA were obtained using a Tapestation 4200 (Agilent) with a D5000 High Sensitivity Assay (Agilent). 5’ single cell RNA-seq libraries were constructed using 50 ng cDNA. After fragmentation, end-repair, A-tailing and size selection purification, the cDNA was ligated with Illumina adaptors and subsequently purified with SPRISelect reagent. Libraries were amplified and barcoded by PCR with dual index oligonucleotides from the Chromium Dual Index TT Set A kit (10X Genomics), for a total of 16 PCR cycles. The final PCR products were subjected to double-sided size selection with SPRISelect beads (0.6x and 0.8x). The library fragment size distributions were analyzed on an Agilent 4200 Tapestation and a Fragment Analyzer, and library yields and concentrations were determined by qPCR using the KAPA Illumina Library Quantification PCR kit (Roche Diagnostics).

Library sequencing was performed at a depth of ∼50,000 paired reads/cell on an Illumina NovaSeq X Plus instrument operated by a commercial provider (Novogene, Inc.), with the following paired-end sequencing settings: Read1, 151 cycles; i7 index, 10 cycles; i5 index, 10 cycles; Read2, 151 cycles.

## Data analysis

Raw sequencing reads were processed using Cell Ranger v8.0.1 to generate gene–cell count matrices. Downstream analysis was performed in R v5.3.1 within RStudio. Filtering was done using perCellQCMetrics and perCellQCFilters functions from the Scater package v1.34.1. Cells with total features or counts more than three median absolute deviations (MADs) below the median were discarded, as were cells with mitochondrial transcript percentages more than three MADs above the median. Identification of mitochondrial genes was done using AnnotationHub v3.14.0. Genes expressed in fewer than two cells were also removed. Doublets were removed using DoubletFinder v2.0.6.

Filtered cells were processed and clustered using the standard Seurat v5 workflow, including normalization, scaling, principal component analysis (PCA), and uniform manifold approximation and projection (UMAP). The number of principal components used for UMAP dimensionality reduction was determined by: the minimum between the point where cumulative variance explained exceeded 90% and the variance contribution of the individual PC was less than 5%, and the point where the drop in variance explained between consecutive PCs was less than 0.1%. Cell types were classified with SingleR v2.8.0 by using the ImmGenData reference from the Celldex package v1.16.0. B cells were subset and clustered using FindNeighbors and FindClusters at resolution 0.3. Low-quality B-cell clusters with median counts below 500 and median features below 200 were removed. The remaining B cells were re-clustered at resolution 0.6, and subtypes were identified using common B-cell markers. Differentially expressed genes were found using FindMarkers using the test.use=wilcox_limma argument. Genes meeting the criteria of p value < 0.05 and log2 fold change > 0.5 were included in the over-representation analysis. Over-representation analysis was done using the clusterProfiler package v.4.14.6 and the gene sets used were from the MsigDB website (https://www.gsea-msigdb.org/gsea/msigdb).

### Immunohistochemistry (IHC)

IHC and IF were performed as previously described ^69^. For IHC, after hydration, antigen retrieval was performed in 10 mM citrate buffer (pH 6.0) by heating the section for 10 min and then allow it to sit for 30 min. The endogenous peroxidase activity was quenched in 3% H_2_O_2_ in MeOH. The section was blocked for 2.5 h with 10% goat serum followed by primary antibody incubation overnight at 4 °C. After washing, the section was incubated with biotinylated secondary antibody for 1 h, then streptavidin-biotin complex (Vectastain Elite ABC kit, Vector Laboratories) was applied for 30 min. Peroxidase substrate solution 3,3′-diaminobenzidine (DAB, Cell Signaling # 8059S) was applied for proper time until desired color reaction is observed. The reaction was terminated by rinsing gently with distilled water. Section was then counterstained with hematoxylin, dehydrated, and coverslipped.

### BCR repertoire profiling

Mouse BCR repertoire profiling from total RNA was performed as described ^70^. In brief, total RNA was isolated from splenocytes and reverse-transcribed into cDNA. Multiplex PCR were used to amplify immunoglobulin (Ig) heavy chain VDJ regions. PCR amplicons were purified using NEBNext Sample Purification Beads and subjected to high-throughput sequencing. Pair-end reads were assembled and UMI-corrected. Bulk IgH repertoire profiling was performed using next-generation sequencing data processed through the Immcantation framework (version 4.4.0). Raw FASTA and IgBLAST output files (.fmt7) were converted into Change-O databases with Change-O v1.3.1 running inside a Conda “immcant” environment (Python 3.10, Bioconda channels). Germline reference sequences for Mus musculus (IGHV, IGHD, IGHJ) were obtained from IMGT/GENE-DB (release 2025-10). IgBLAST v1.22.0 was used for V(D)J alignment with the corresponding IMGT germline database. Productive IGH reads were identified (locus = IGH, productive = True), and clonal clustering was performed using a Hamming-distance model (--model ham, --dist 0.15, --norm len, --act set) to group sequences sharing identical V and J genes and CDR3 length within a normalized junction distance ≤15% normalized Hamming distance across the junction region to account for somatic hypermutation. Downstream tabulation and visualization were done in Python 3.10 using pandas v2.2 and matplotlib v3.8. Constant-region (C-gene) assignments (c_call) were derived from IgBLAST to determine isotype usage. Clonal frequencies were expressed as the fraction of total unique molecular identifier (UMI)-normalized sequences per sample ^71^.

## Data and materials availability

All data are available in the main text or the supplementary materials. The CRISPR/Cas9 screen in Ba/F3 cells was deposited into the NCBI Gene Expression Omnibus (GEO) with the accession number GSE341452. Spleen tissue bulk RNA-seq was deposited into the GEO with the accession number GSE341453. The scRNA-seq comparing Cdkn2a-/- and DKO mice is GSE338997. The mass spectrometry proteomics data have been deposited to the ProteomeXchange Consortium via the PRIDE partner repository with the dataset identifier PXD081511 and 10.6019/PXD081511.

## Statistical analysis

Data was analyzed using GraphPad Prism 8 software. Data were presented as mean ± SD. Unpaired Student’s t test was used to compare between two groups, differences among more than two groups using one-way ANOVA. Two-way ANOVA with Sidak’s multiple comparisons test was used to compare the effects of two independent variables (factors) on a dependent variable among multiple groups. Log-rank (Mantel-Cox) test was used to analyze statistical significance in the Kaplan-Meier survival plots. Statistical significance was judged when p<0.05.

## Acknowledgements

Single cell RNAseq experiments were conducted by the Northport VAMC/Stony Brook University Single Cell and Spatial Multiomics Facility directed by Barbara Rosati. The study was supported by the NIH (R01CA129536, R01CA224550, and R01CA286043 to WXZ; R01CA293055 to PX; S10OD034300 and S10OD038256 to HL).

## Author Contributions

MUT performed experiments and wrote the paper. NS, JS, JJ, RL, KL, JY, HS, and TL performed additional experiments. MCK and GC performed scRNA-seq and data analysis. BV and JW performed computational analysis. SKB, HL, CH, RZL, YLW, and JV contributed methodology and experimental resources. YS performed histopathological characterization and statistical analysis. FB performed patient data analysis. PX and WXZ conceptualized the project and wrote the paper. All authors have read and approved the final version of the manuscript.

## Supplemental Figures

**Suppl. Fig. S1.**
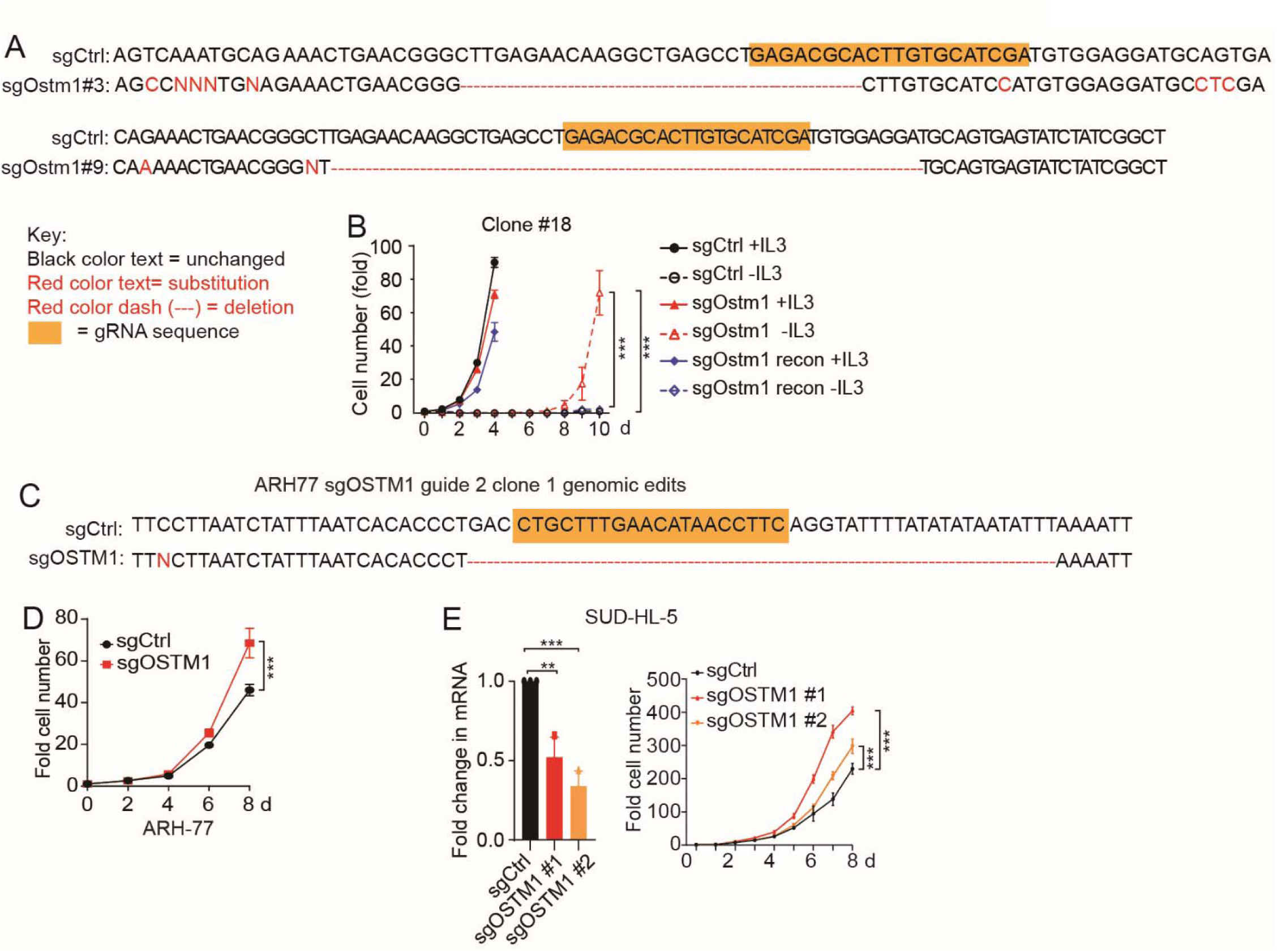
(related to Fig. 1). **(A)** Ostm1 was silenced in Ba/F3 cells using two independent sgRNAs. Successful knockout of Ostm1 was determined by genomic sequencing, which shows frame-shifting deletion, point mutations, and large-size deletion. **(B)** Human OSTM1-Flag was reconstituted into sg*Ostm1* Ba/F3 cells, which led to the loss of IL3-independence of the sg*Ostm1* cells. **(C)** OSTM1 was silenced in ARH-77 cells using sgRNA. Successful knockout of OSTM1 was determined by genomic sequencing. **(C)** Silencing OSTM1 in ARH-77 cells promoted cell proliferation. **(D)** OSTM1 was silenced in SU-DHL-5 cells with two sgRNAs. Successful silencing was achieved indicated by decreased mRNA levels, which led to increased cell proliferation. Statistical analysis in panels **B**, **D**, and **E** was done using multiple t-test, one per row. For panel **E**, one way ANOVA was used. **p<0.01; ***p<0.001.

**Suppl. Fig. S2.**
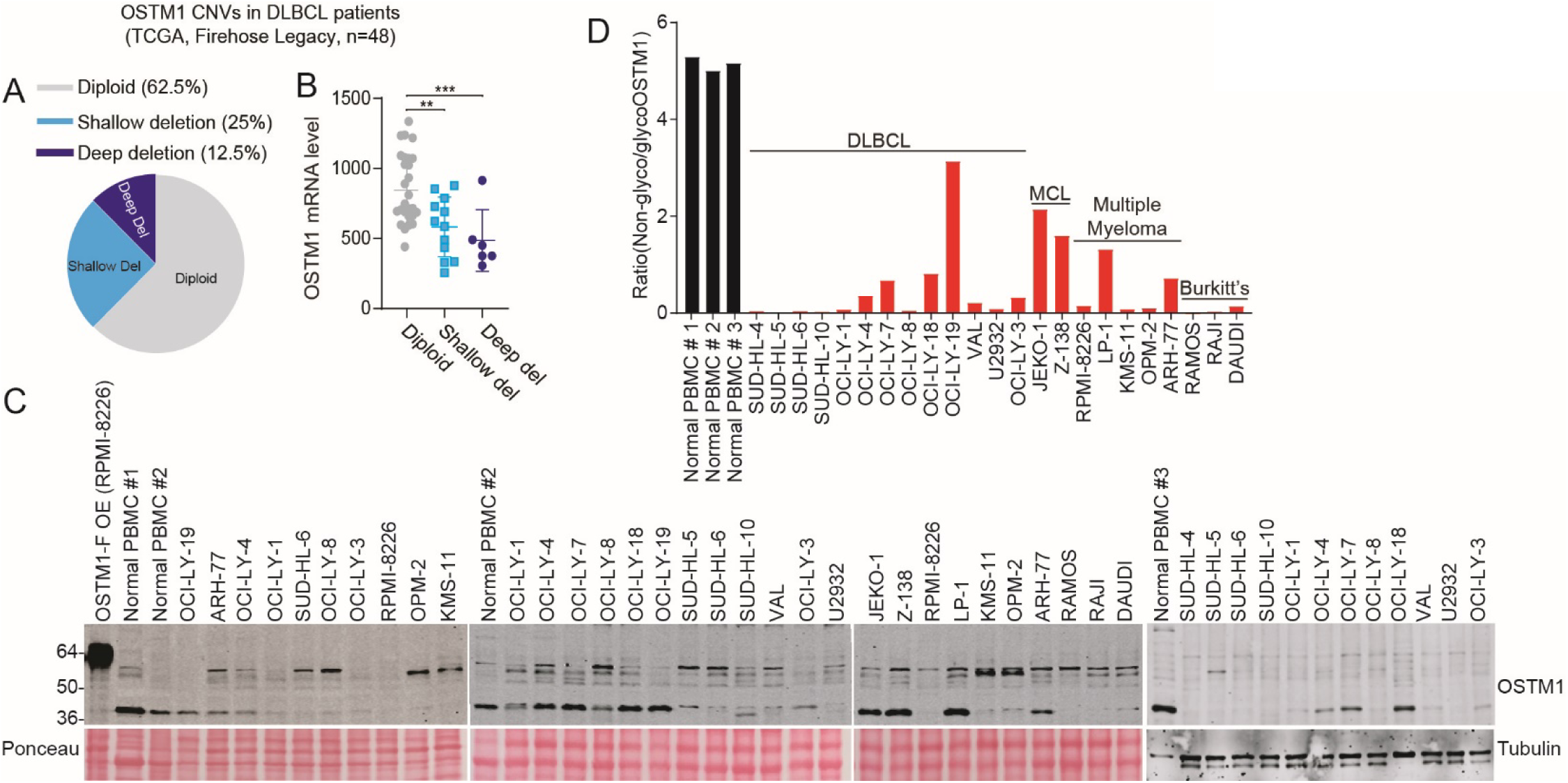
(related to Fig. 2). **(A)** *OSTM1* copy number variations (CNVs) were available in DLBCL patients in TCGA Firehose Legacy. While 25% patients had shallow deletion, 12.5% had deep deletion. **(B)** *OSTM1* deletions correlated with decreased mRNA expression in the same patients. **(C)** RPMI-8226 cell line stably expressing OSTM1-Flag, PBMC isolated from three healthy donors, and indicated BCL cell lines were probed for ectopically expressed or endogenous OSTM1. **(D)** ImageJ quantification of the ratio between non-glycosylated (∼37 kDa) and glycosylated (∼60 kDa) OSTM1.

**Suppl. Fig. S3.**
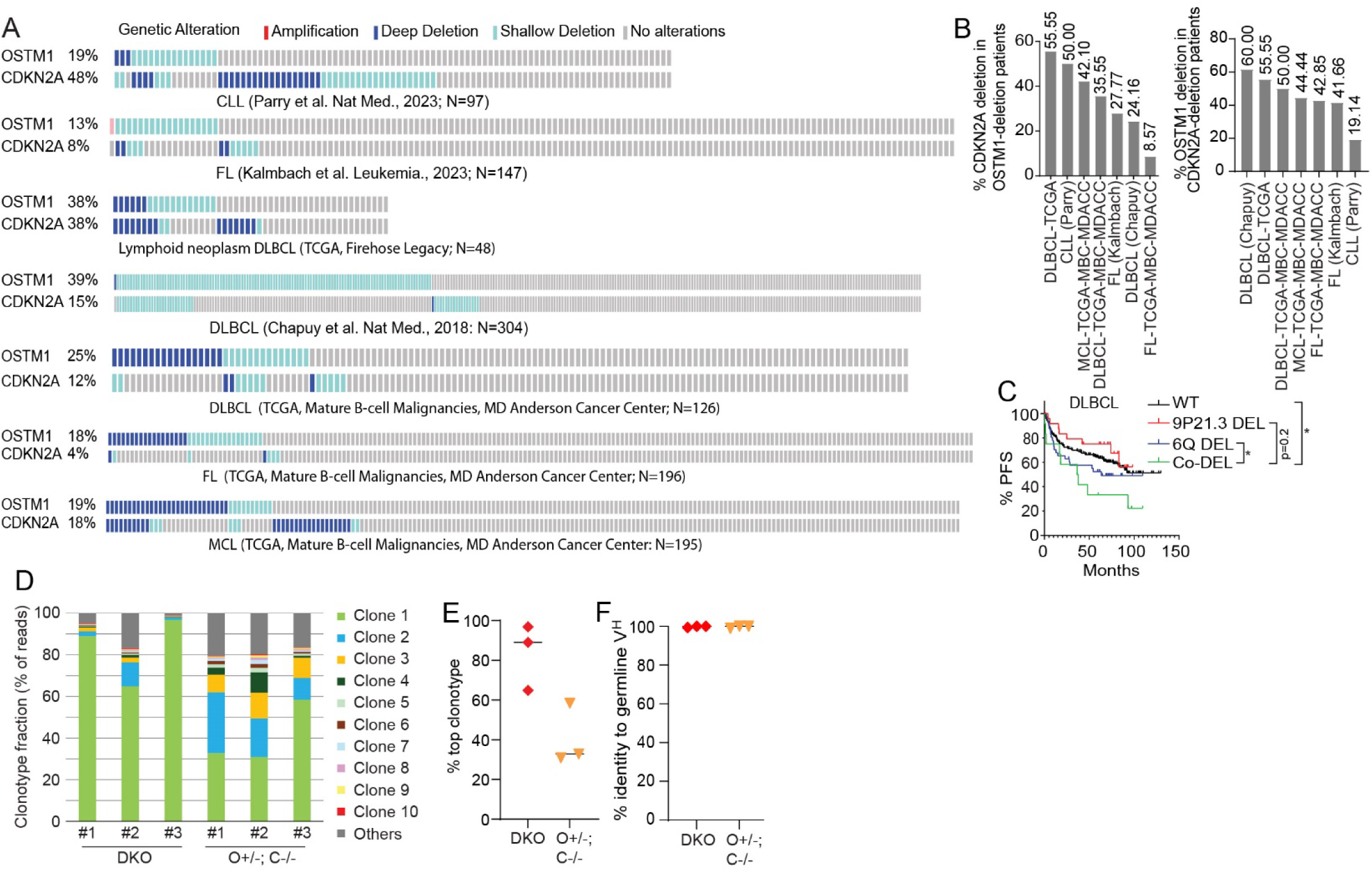
(related to Fig. 3). **(A)** Oncoprint visualization of *OSTM1* and *CDKN2A* co-deletions across different BCLs as reported in the indicated studies. **(B)** Frequencies of *CDKN2A* deletions among patients with *OSTM1* deletions across BCLs (left), and the frequencies of *OSTM1* deletion among patients with *CDKN2A* deletions (right), in the indicated studies. **(C)** Progression-free survival of DLBCL patients ^20^ stratified by the presence of chr. 6q deletion (*OSTM1* DEL), 9p21.3 deletion (*CDKN2A* DEL), both (*Co-DEL*), or neither (WT). Patients with co-deletions had worse survival. (**D**) Frequencies of individual clonotypes obtained from Igh V(D)J sequencing of mouse spleen samples. Each clonotype is color-coded according to its rank of prevalence within the corresponding sample. (**E**) Comparison of the frequency of the most expanded clonotype across samples, grouped by genotype. (**F**) Percent identity between BCL Igh V(D)J sequences and their corresponding germline sequences, grouped by genotype.

**Suppl. Fig. S4.**
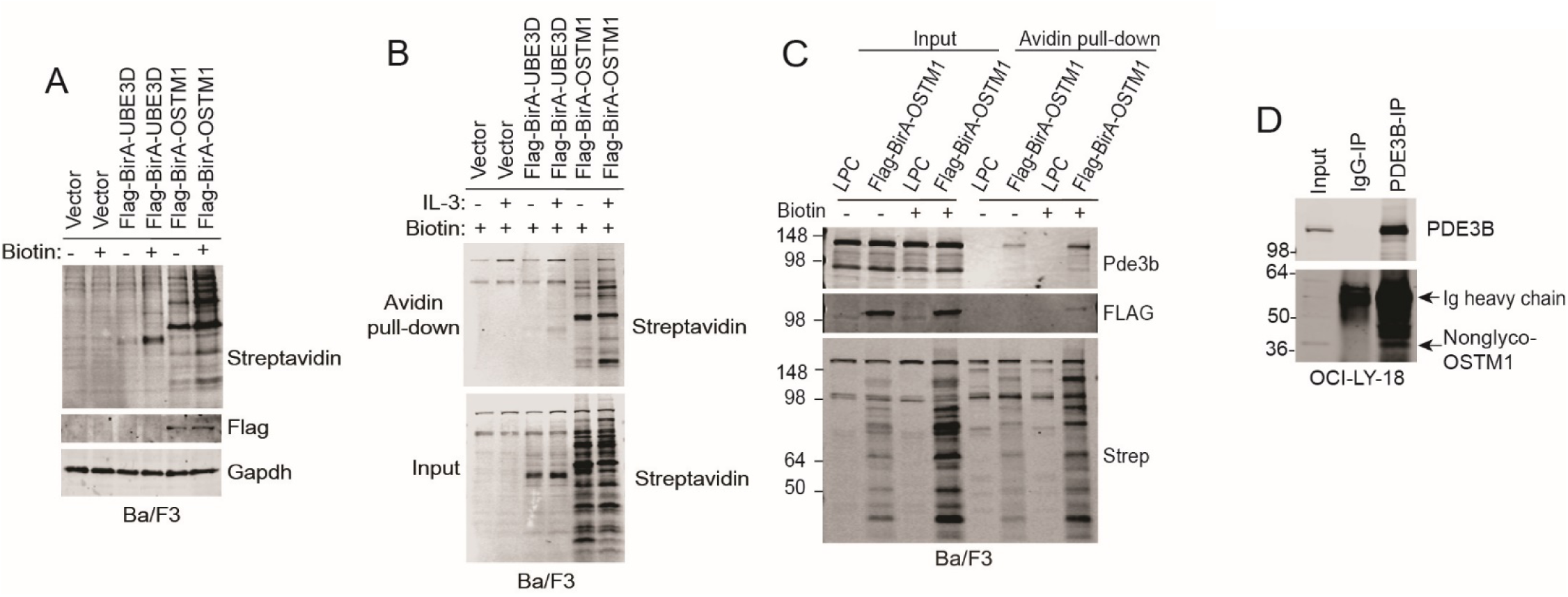
(related to Fig. 4). **(A)** Flag-BirA-OSTM1 was stably expressed in Ba/F3 cells. 50 µM biotin was added to the cells for 24 h. Cells were lysed in BioID lysis buffer and the BirA activity was confirmed by HRP-conjugated streptavidin in IB. Successful expression of Flag-BirA-OSTM1 was confirmed by Flag IB. The lanes showing Flag-BirA-UBE3D were for a different project. **(B)** Ba/F3 cells stably expressing Flag-BirA-OSTM1 were cultured in the absence or presence of IL3. Biotin was added to the cell lysates. Avidin pull-down was performed and probed for streptavidin by IB. Biotinolyted proteins were detected. **(C)** Avidin beads pull-down was performed on Ba/F3 cells expressing Flag-BirA-OSTM1, followed by IB with the indicated antibodies. Note that Pde3b was detected to interact with Flag-BirA-OSTM1. **(D)** OSTM1 interacts with PDE3B at the endogenous level. Endogenous PDE3B was immunoprecipitated from OCI-LY-18 lysates with PDE3B antibody or IgG as isotype control. Subsequently, IB was performed using PDE3B and OSTM1 antibodies.

**Suppl. Fig. S5.**
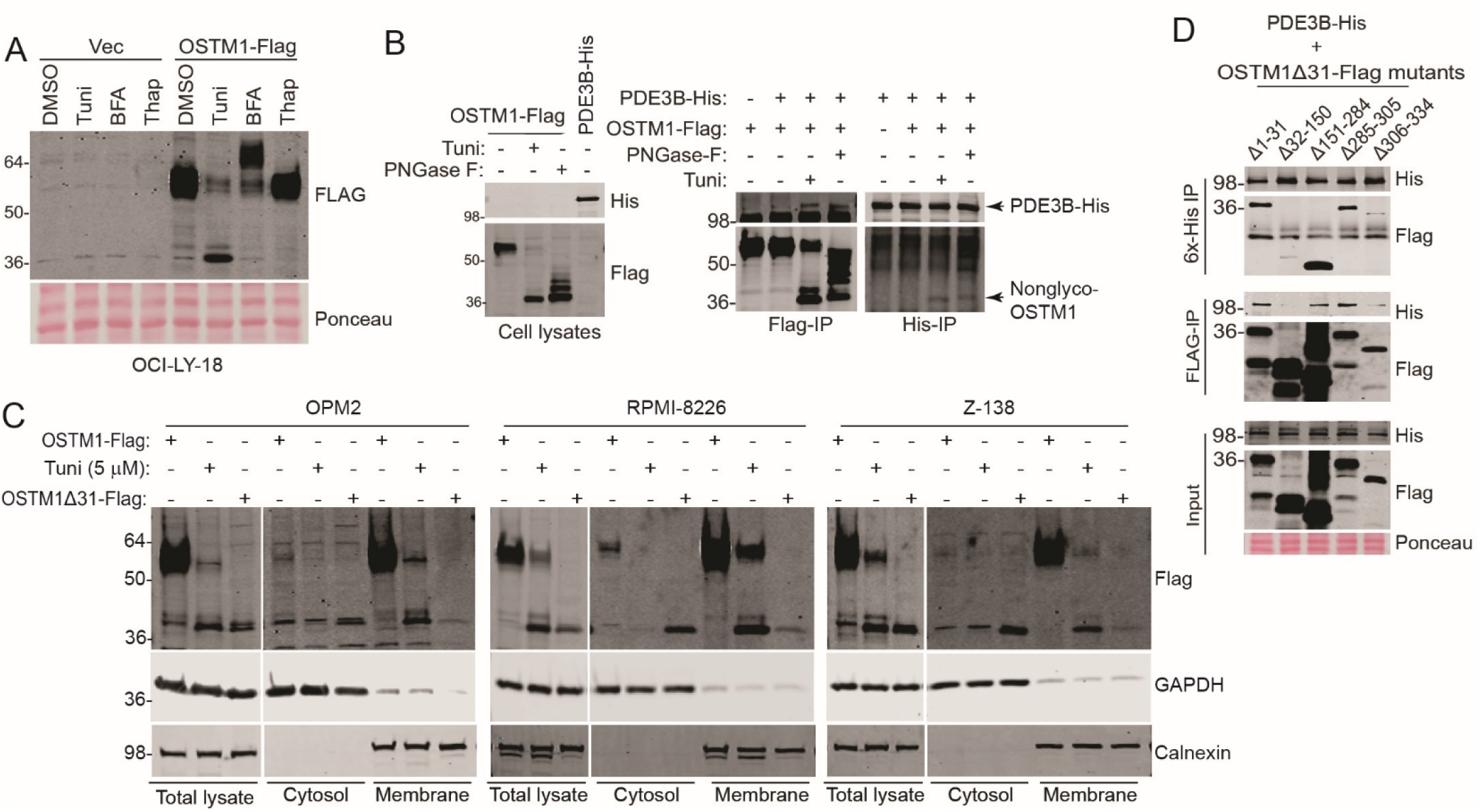
(related to Fig. 5). **(A)** OCI-LY-18 cells stably expressing OSTM1-Flag were treated with tunicamycin (Tuni, 5 µg/ml), brefeldin A (BFA, 3 µg/ml), and thapsigargin (Thap, 1 µM). Note that only Tuni led to the accumulation of the ∼37 kD band of OSTM1. **(B)** HEK293T cells stably expressing OSTM1-Flag were treated with tunicamycin (5 µg/ml, 24 h) or PNGase-F (200 μg lysate protein/1,000 units, 4 h), and cell lysates were made. Cell lysates were also made from HEK293T cells stably expressing PDE3B. Indicated cell lysates were mixed and Flag or His antibody IP were performed. Note that tunicamycin and PNGase-F treatments led to de-glycosylated OSTM1, which showed stronger interactions with PDE3B than the glycosylated OSTM1 in untreated cell lysates. **(C)** OSTM1-Flag or OSTM1Δ31-Flag was stably expressed in the indicated BCL cell lines. Cells expressing OSTM1-Flag were treated with tunicamycin (5 µg/ml, 24 h). Cytosol/membrane fractionation was performed, and the indicated proteins were detected. Note that tunicamycin leads to de-glycosylation of OSTM1, and the signal peptide (SP)-less OSTM1Δ31 is predominantly localized in the cytosol. **(D)** PDE3B-His and indicated mutants of OSTM1Δ31-Flag were co-transfected into HEK293T cells. Reciprocal IP with His or Flag antibody was performed. Note that the CTD-less (Δ306-334) mutant lost the interaction with PDE3B.

**Suppl. Fig. S6.**
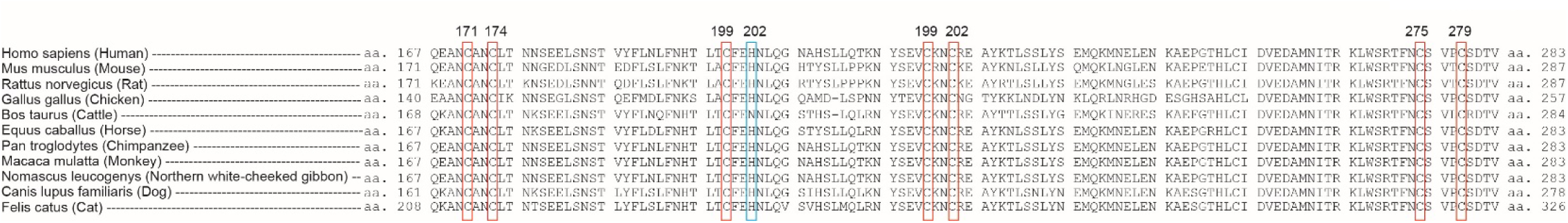
(related to Fig. 6). Sequence aligment of the RING-like domains in OSTM1 among various species. The putative RING-like domains of OSTM1 from various species are aligned. The Cys-Cys and Cys-His zinc-coordination clusters are highlighted.

**Suppl. Fig. S7.**
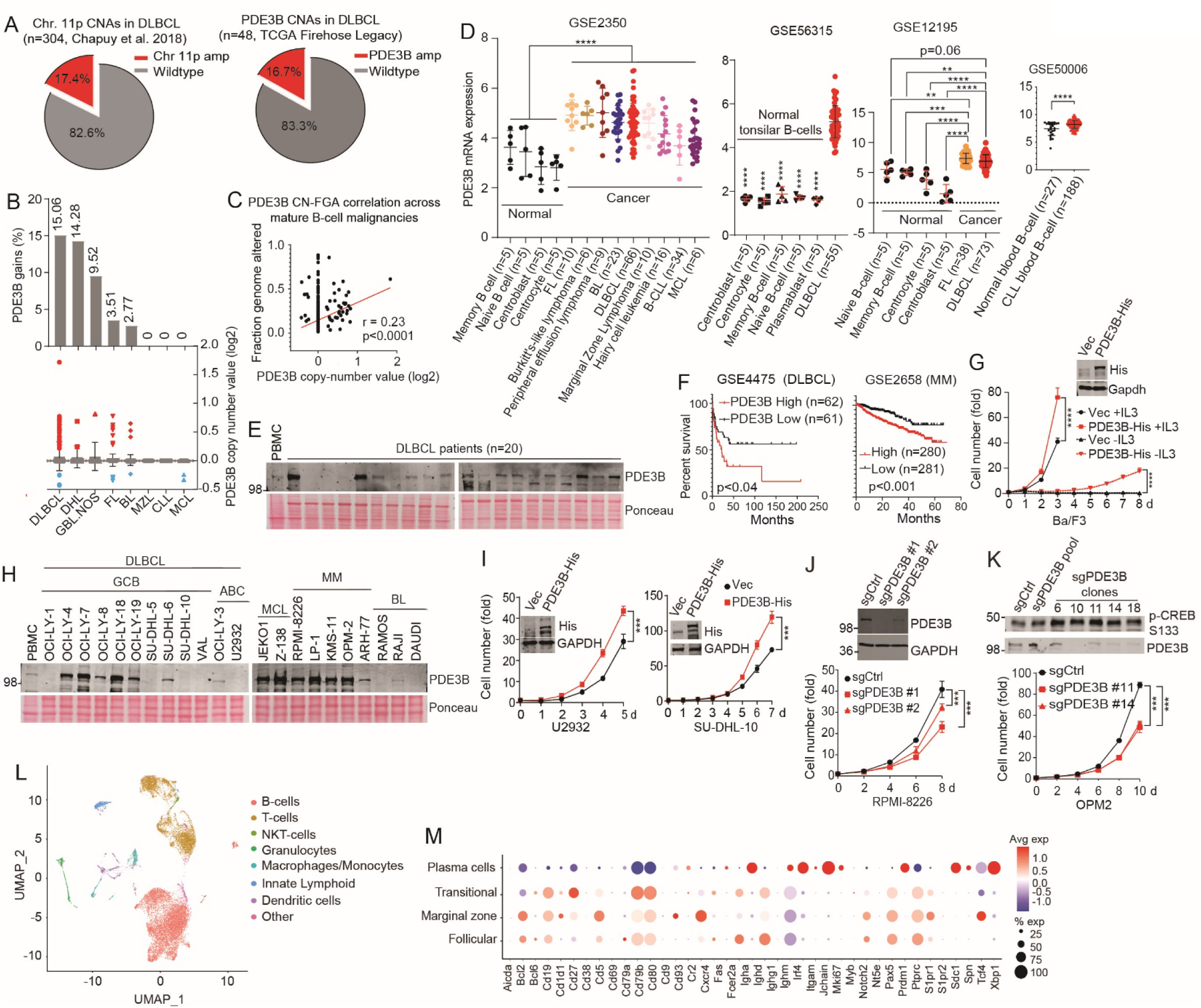
(related to Fig. 7). **(A)** Left: Pie chart showing the percentage of DLBCL patients^20^ with chr. 11p amplification (17.4%). Right: Pie chart showing the fraction of DLBCL patients (n = 48, TCGA Firehose Legacy) with PDE3B amplification (16.7%). **(B)** Percentage of patients with PDE3B gains across B-cell malignancies (TCGA, MDACC dataset), with individual log2 copy-number values overlaid. **(C)** Spearman correlation between PDE3B log2 copy-number value and fraction genome altered (FGA) in mature B-cell malignancies. **(D)** PDE3B expression across indicated BCL cohorts, comparing normal B cells with BCLs; P values were calculated by grouping all normal samples and cancer samples separately and comparing the two groups using Student’s t-test. **(E)** DLBCL patient samples were obtained from the Rutgers Cancer Institute. The tumor tissue lysates and PBMC from a healthy donor were probed for PDE3B expression. **(F)** Kaplan-Meier survival curves in DLBCL (GSE4475) and multiple myeloma (GSE2658) patients showing reduced survival in patients with high PDE3B expression, stratified by median expression. **(G)** PDE3B-His was stably expressed in Ba/F3 cells. Cells were cultured in the presence or absence of IL3, and cell growth was determined by counting cell numbers. ****p<0.0001. **(H)** PDE3B was detected by IB across cell lines from different BCL subtypes. **(I)** PDE3B-His was stably expressed in indicated BCL cell lines, and cell proliferation was measured. ***p<0.001. **(J and K)** PDE3B was silenced with 2 sgRNAs in RPMI-8226 and OPM2 cell lines. Cell proliferation was measured. P-value calculation was done using multiple t-test comparisons, one per row ***p<0.001; ****p<0.0001. **(L and M)** Spleens from two-month-old C^-/-^ and DKO mice were harvested for scRNA-seq. **(L)** Pan-cell-type UMAP embeddings with cells from both C^-/-^ and DKO mice combined (n=10,000 cells per mouse). **(M)** Dot-plot of B-cell markers stratified by sub-population. Color represents average log fold expression.

## References

1. Merup, M., Moreno, T.C., Heyman, M., Ronnberg, K., Grander, D., Detlofsson, R., Rasool, O., Liu, Y., Soderhall, S., Juliusson, G., Gahrton, G. & Einhorn, S. 6q deletions in acute lymphoblastic leukemia and non-Hodgkin’s lymphomas. Blood 91, 3397–3400 (1998).

2. Remke, M., Pfister, S., Kox, C., Toedt, G., Becker, N., Benner, A., Werft, W., Breit, S., Liu, S., Engel, F., Wittmann, A., Zimmermann, M., Stanulla, M., Schrappe, M., Ludwig, W.D., Bartram, C.R., Radlwimmer, B., Muckenthaler, M.U., Lichter, P. & Kulozik, A.E. High-resolution genomic profiling of childhood T-ALL reveals frequent copy-number alterations affecting the TGF-beta and PI3K-AKT pathways and deletions at 6q15-16.1 as a genomic marker for unfavorable early treatment response. Blood 114, 1053–1062 (2009).

3. Bonn, B.R., Rohde, M., Zimmermann, M., Krieger, D., Oschlies, I., Niggli, F., Wrobel, G., Attarbaschi, A., Escherich, G., Klapper, W., Reiter, A. & Burkhardt, B. Incidence and prognostic relevance of genetic variations in T-cell lymphoblastic lymphoma in childhood and adolescence. Blood 121, 3153–3160 (2013).

4. Jarosova, M., Hruba, M., Oltova, A., Plevova, K., Kruzova, L., Kriegova, E., Fillerova, R., Koritakova, E., Doubek, M., Lysak, D., Prochazka, V., Mraz, M., Indrak, K. & Papajik, T. Chromosome 6q deletion correlates with poor prognosis and low relative expression of FOXO3 in chronic lymphocytic leukemia patients. Am J Hematol 92, E604–E607 (2017).

5. Siu, L.L., Chan, V., Chan, J.K., Wong, K.F., Liang, R. & Kwong, Y.L. Consistent patterns of allelic loss in natural killer cell lymphoma. Am J Pathol 157, 1803–1809 (2000).PMC1885756.

6. Palacios, R. & Steinmetz, M. Il-3-dependent mouse clones that express B-220 surface antigen, contain Ig genes in germ-line configuration, and generate B lymphocytes in vivo. Cell 41, 727–734 (1985).

7. Daley, G.Q. & Baltimore, D. Transformation of an interleukin 3-dependent hematopoietic cell line by the chronic myelogenous leukemia-specific P210bcr/abl protein. Proc Natl Acad Sci U S A 85, 9312–9316 (1988).PMC282729.

8. Bai, R.Y., Dieter, P., Peschel, C., Morris, S.W. & Duyster, J. Nucleophosmin-anaplastic lymphoma kinase of large-cell anaplastic lymphoma is a constitutively active tyrosine kinase that utilizes phospholipase C-gamma to mediate its mitogenicity. Mol Cell Biol 18, 6951–6961 (1998).PMC109278.

9. Kelly, L.M., Liu, Q., Kutok, J.L., Williams, I.R., Boulton, C.L. & Gilliland, D.G. FLT3 internal tandem duplication mutations associated with human acute myeloid leukemias induce myeloproliferative disease in a murine bone marrow transplant model. Blood 99, 310–318 (2002).

10. Carroll, M., Tomasson, M.H., Barker, G.F., Golub, T.R. & Gilliland, D.G. The TEL/platelet-derived growth factor beta receptor (PDGF beta R) fusion in chronic myelomonocytic leukemia is a transforming protein that self-associates and activates PDGF beta R kinase-dependent signaling pathways. Proc Natl Acad Sci U S A 93, 14845–14850 (1996).PMC26224.

11. Lacronique, V., Boureux, A., Valle, V.D., Poirel, H., Quang, C.T., Mauchauffe, M., Berthou, C., Lessard, M., Berger, R., Ghysdael, J. & Bernard, O.A. A TEL-JAK2 fusion protein with constitutive kinase activity in human leukemia. Science 278, 1309–1312 (1997).

12. Chalhoub, N., Benachenhou, N., Rajapurohitam, V., Pata, M., Ferron, M., Frattini, A., Villa, A. & Vacher, J. Grey-lethal mutation induces severe malignant autosomal recessive osteopetrosis in mouse and human. Nat Med 9, 399–406 (2003).

13. Pangrazio, A., Poliani, P.L., Megarbane, A., Lefranc, G., Lanino, E., Di Rocco, M., Rucci, F., Lucchini, F., Ravanini, M., Facchetti, F., Abinun, M., Vezzoni, P., Villa, A. & Frattini, A. Mutations in OSTM1 (grey lethal) define a particularly severe form of autosomal recessive osteopetrosis with neural involvement. J Bone Miner Res 21, 1098–1105 (2006).

14. Heraud, C., Griffiths, A., Pandruvada, S.N., Kilimann, M.W., Pata, M. & Vacher, J. Severe neurodegeneration with impaired autophagy mechanism triggered by ostm1 deficiency. J Biol Chem 289, 13912–13925 (2014).PMC4022863.

15. Fischer, T., De Vries, L., Meerloo, T. & Farquhar, M.G. Promotion of G alpha i3 subunit down-regulation by GIPN, a putative E3 ubiquitin ligase that interacts with RGS-GAIP. Proc Natl Acad Sci U S A 100, 8270–8275 (2003).PMC166218.

16. Pandruvada, S.N., Beauregard, J., Benjannet, S., Pata, M., Lazure, C., Seidah, N.G. & Vacher, J. Role of Ostm1 Cytosolic Complex with Kinesin 5B in Intracellular Dispersion and Trafficking. Mol Cell Biol 36, 507–521 (2016).PMC4719428.

17. Pata, M., Yousefi Behzadi, P. & Vacher, J. Expression pattern of the V5-Ostm1 protein in bacterial artificial chromosome transgenic mice. Genesis 59, e23409 (2021).

18. Burkhardt, B., Michgehl, U., Rohde, J., Erdmann, T., Berning, P., Reutter, K., Rohde, M., Borkhardt, A., Burmeister, T., Dave, S., Tzankov, A., Dugas, M., Sandmann, S., Fend, F., Finger, J., Mueller, S., Gokbuget, N., Haferlach, T., Kern, W., Hartmann, W., Klapper, W., Oschlies, I., Richter, J., Kontny, U., Lutz, M., Maecker-Kolhoff, B., Ott, G., Rosenwald, A., Siebert, R., von Stackelberg, A., Strahm, B., Woessmann, W., Zimmermann, M., Zapukhlyak, M., Grau, M. & Lenz, G. Clinical relevance of molecular characteristics in Burkitt lymphoma differs according to age. Nat Commun 13, 3881 (2022).PMC9259584 Leukamielabor GmbH. The remaining authors declare no competing interests.

19. Parry, E.M., Leshchiner, I., Guieze, R., Johnson, C., Tausch, E., Parikh, S.A., Lemvigh, C., Broseus, J., Hergalant, S., Messer, C., Utro, F., Levovitz, C., Rhrissorrakrai, K., Li, L., Rosebrock, D., Yin, S., Deng, S., Slowik, K., Jacobs, R., Huang, T., Li, S., Fell, G., Redd, R., Lin, Z., Knisbacher, B.A., Livitz, D., Schneider, C., Ruthen, N., Elagina, L., Taylor-Weiner, A., Persaud, B., Martinez, A., Fernandes, S.M., Purroy, N., Anandappa, A.J., Ma, J., Hess, J., Rassenti, L.Z., Kipps, T.J., Jain, N., Wierda, W., Cymbalista, F., Feugier, P., Kay, N.E., Livak, K.J., Danysh, B.P., Stewart, C., Neuberg, D., Davids, M.S., Brown, J.R., Parida, L., Stilgenbauer, S., Getz, G. & Wu, C.J. Evolutionary history of transformation from chronic lymphocytic leukemia to Richter syndrome. Nat Med 29, 158–169 (2023).PMC10155825.

20. Chapuy, B., Stewart, C., Dunford, A.J., Kim, J., Kamburov, A., Redd, R.A., Lawrence, M.S., Roemer, M.G.M., Li, A.J., Ziepert, M., Staiger, A.M., Wala, J.A., Ducar, M.D., Leshchiner, I., Rheinbay, E., Taylor-Weiner, A., Coughlin, C.A., Hess, J.M., Pedamallu, C.S., Livitz, D., Rosebrock, D., Rosenberg, M., Tracy, A.A., Horn, H., van Hummelen, P., Feldman, A.L., Link, B.K., Novak, A.J., Cerhan, J.R., Habermann, T.M., Siebert, R., Rosenwald, A., Thorner, A.R., Meyerson, M.L., Golub, T.R., Beroukhim, R., Wulf, G.G., Ott, G., Rodig, S.J., Monti, S., Neuberg, D.S., Loeffler, M., Pfreundschuh, M., Trumper, L., Getz, G. & Shipp, M.A. Molecular subtypes of diffuse large B cell lymphoma are associated with distinct pathogenic mechanisms and outcomes. Nat Med 24, 679–690 (2018).PMC6613387.

21. Frontzek, F., Staiger, A.M., Wullenkord, R., Grau, M., Zapukhlyak, M., Kurz, K.S., Horn, H., Erdmann, T., Fend, F., Richter, J., Klapper, W., Lenz, P., Hailfinger, S., Tasidou, A., Trautmann, M., Hartmann, W., Rosenwald, A., Quintanilla-Martinez, L., Ott, G., Anagnostopoulos, I. & Lenz, G. Molecular profiling of EBV associated diffuse large B-cell lymphoma. Leukemia 37, 670–679 (2023).PMC9991915

22. Kalmbach, S., Grau, M., Zapukhlyak, M., Leich, E., Jurinovic, V., Hoster, E., Staiger, A.M., Kurz, K.S., Weigert, O., Gaitzsch, E., Passerini, V., Engelhard, M., Herfarth, K., Beiske, K., Micci, F., Moller, P., Bernd, H.W., Feller, A.C., Klapper, W., Stein, H., Hansmann, M.L., Hartmann, S., Dreyling, M., Holte, H., Lenz, G., Rosenwald, A., Ott, G., Horn, H. & German Lymphoma, A. Novel insights into the pathogenesis of follicular lymphoma by molecular profiling of localized and systemic disease forms. Leukemia 37, 2058–2065 (2023).PMC10539171.

23. Hoang, P.H., Cornish, A.J., Sherborne, A.L., Chubb, D., Kimber, S., Jackson, G., Morgan, G.J., Cook, G., Kinnersley, B., Kaiser, M. & Houlston, R.S. An enhanced genetic model of relapsed IGH-translocated multiple myeloma evolutionary dynamics. Blood Cancer J 10, 101 (2020).PMC7560599.

24. Oricchio, E., Katanayeva, N., Donaldson, M.C., Sungalee, S., Pasion, J.P., Beguelin, W., Battistello, E., Sanghvi, V.R., Jiang, M., Jiang, Y., Teater, M., Parmigiani, A., Budanov, A.V., Chan, F.C., Shah, S.P., Kridel, R., Melnick, A.M., Ciriello, G. & Wendel, H.G. Genetic and epigenetic inactivation of SESTRIN1 controls mTORC1 and response to EZH2 inhibition in follicular lymphoma. Sci Transl Med 9 (2017).PMC5559734.

25. Lange, P.F., Wartosch, L., Jentsch, T.J. & Fuhrmann, J.C. ClC-7 requires Ostm1 as a beta-subunit to support bone resorption and lysosomal function. Nature 440, 220–223 (2006).

26. Pata, M. & Vacher, J. Ostm1 Bifunctional Roles in Osteoclast Maturation: Insights From a Mouse Model Mimicking a Human OSTM1 Mutation. J Bone Miner Res 33, 888–898 (2018).

27. Rickert, R.C., Roes, J. & Rajewsky, K. B lymphocyte-specific, Cre-mediated mutagenesis in mice. Nucleic Acids Res 25, 1317–1318 (1997).PMC146582.

28. Sewastianik, T., Jiang, M., Sukhdeo, K., Patel, S.S., Roberts, K., Kang, Y., Alduaij, A., Dennis, P.S., Lawney, B., Liu, R., Song, Z., Xiong, J., Zhang, Y., Lemieux, M.E., Pinkus, G.S., Rich, J.N., Weinstock, D.M., Mullighan, C.G., Sharpless, N.E. & Carrasco, R.D. Constitutive Ras signaling and Ink4a/Arf inactivation cooperate during the development of B-ALL in mice. Blood Adv 1, 2361–2374 (2017).PMC5729631 interests.

29. Vacher, J., Bruccoleri, M. & Pata, M. Ostm1 from Mouse to Human: Insights into Osteoclast Maturation. Int J Mol Sci 21 (2020).PMC7460669.

30. Degerman, E., Belfrage, P. & Manganiello, V.C. Structure, localization, and regulation of cGMP-inhibited phosphodiesterase (PDE3). J Biol Chem 272, 6823–6826 (1997).

31. Wechsler, J., Choi, Y.H., Krall, J., Ahmad, F., Manganiello, V.C. & Movsesian, M.A. Isoforms of cyclic nucleotide phosphodiesterase PDE3A in cardiac myocytes. J Biol Chem 277, 38072–38078 (2002).

32. Schrecker, M., Korobenko, J. & Hite, R.K. Cryo-EM structure of the lysosomal chloride-proton exchanger CLC-7 in complex with OSTM1. Elife 9 (2020).PMC7440919.

33. Zhang, S., Liu, Y., Zhang, B., Zhou, J., Li, T., Liu, Z., Li, Y. & Yang, M. Molecular insights into the human CLC-7/Ostm1 transporter. Sci Adv 6, eabb4747 (2020).PMC7423370.

34. Degerman, E., Ahmad, F., Chung, Y.W., Guirguis, E., Omar, B., Stenson, L. & Manganiello, V. From PDE3B to the regulation of energy homeostasis. Curr Opin Pharmacol 11, 676–682 (2011).PMC3225700.

35. Berger, K., Lindh, R., Wierup, N., Zmuda-Trzebiatowska, E., Lindqvist, A., Manganiello, V.C. & Degerman, E. Phosphodiesterase 3B is localized in caveolae and smooth ER in mouse hepatocytes and is important in the regulation of glucose and lipid metabolism. PLoS One 4, e4671 (2009).PMC2650791.

36. Lugnier, C. Cyclic nucleotide phosphodiesterase (PDE) superfamily: a new target for the development of specific therapeutic agents. Pharmacol Ther 109, 366–398 (2006).

37. Garcia-Barcena, C., Osinalde, N., Ramirez, J. & Mayor, U. How to Inactivate Human Ubiquitin E3 Ligases by Mutation. Front Cell Dev Biol 8, 39 (2020).PMC7010608.

38. Ahmad, F., Murata, T., Shimizu, K., Degerman, E., Maurice, D. & Manganiello, V. Cyclic nucleotide phosphodiesterases: important signaling modulators and therapeutic targets. Oral Dis 21, e25–50 (2015).PMC4275405.

39. Dou, A.X. & Wang, X. Cyclic adenosine monophosphate signal pathway in targeted therapy of lymphoma. Chin Med J (Engl) 123, 95–99 (2010).

40. Lerner, A., Kim, D.H. & Lee, R. The cAMP signaling pathway as a therapeutic target in lymphoid malignancies. Leuk Lymphoma 37, 39–51 (2000).

41. Lerner, A. & Epstein, P.M. Cyclic nucleotide phosphodiesterases as targets for treatment of haematological malignancies. Biochem J 393, 21–41 (2006).PMC1383661.

42. Murray, F. & Insel, P.A. Targeting cAMP in chronic lymphocytic leukemia: a pathway-dependent approach for the treatment of leukemia and lymphoma. Expert Opin Ther Targets 17, 937–949 (2013).

43. Pasqualucci, L., Dominguez-Sola, D., Chiarenza, A., Fabbri, G., Grunn, A., Trifonov, V., Kasper, L.H., Lerach, S., Tang, H., Ma, J., Rossi, D., Chadburn, A., Murty, V.V., Mullighan, C.G., Gaidano, G., Rabadan, R., Brindle, P.K. & Dalla-Favera, R. Inactivating mutations of acetyltransferase genes in B-cell lymphoma. Nature 471, 189–195 (2011).PMC3271441.

44. Zhang, J., Vlasevska, S., Wells, V.A., Nataraj, S., Holmes, A.B., Duval, R., Meyer, S.N., Mo, T., Basso, K., Brindle, P.K., Hussein, S., Dalla-Favera, R. & Pasqualucci, L. The CREBBP Acetyltransferase Is a Haploinsufficient Tumor Suppressor in B-cell Lymphoma. Cancer Discov 7, 322–337 (2017).PMC5386396.

45. Zhang, L., Zambon, A.C., Vranizan, K., Pothula, K., Conklin, B.R. & Insel, P.A. Gene expression signatures of cAMP/protein kinase A (PKA)-promoted, mitochondrial-dependent apoptosis. Comparative analysis of wild-type and cAMP-deathless S49 lymphoma cells. J Biol Chem 283, 4304–4313 (2008).PMC3882191.

46. Mamani-Matsuda, M., Moynet, D., Molimard, M., Ferry-Dumazet, H., Marit, G., Reiffers, J. & Mossalayi, M.D. Long-acting beta2-adrenergic formoterol and salmeterol induce the apoptosis of B-chronic lymphocytic leukaemia cells. Br J Haematol 124, 141–150 (2004).

47. Zhang, H., Kong, Q., Wang, J., Jiang, Y. & Hua, H. Complex roles of cAMP-PKA-CREB signaling in cancer. Exp Hematol Oncol 9, 32 (2020).PMC7684908.

48. Deng, J., Pan, T., Liu, Z., McCarthy, C., Vicencio, J.M., Cao, L., Alfano, G., Suwaidan, A.A., Yin, M., Beatson, R. & Ng, T. The role of TXNIP in cancer: a fine balance between redox, metabolic, and immunological tumor control. British journal of cancer 129, 1877–1892 (2023).PMC10703902.

49. Ci, W., Polo, J.M. & Melnick, A. B-cell lymphoma 6 and the molecular pathogenesis of diffuse large B-cell lymphoma. Current opinion in hematology 15, 381–390 (2008).PMC2748732.

50. Bologna, C., Buonincontri, R., Serra, S., Vaisitti, T., Audrito, V., Brusa, D., Pagnani, A., Coscia, M., D’Arena, G., Mereu, E., Piva, R., Furman, R.R., Rossi, D., Gaidano, G., Terhorst, C. & Deaglio, S. SLAMF1 regulation of chemotaxis and autophagy determines CLL patient response. The Journal of clinical investigation 126, 181–194 (2016).PMC4701571.

51. Schuhmacher, B., Bein, J., Rausch, T., Benes, V., Tousseyn, T., Vornanen, M., Ponzoni, M., Thurner, L., Gascoyne, R., Steidl, C., Kuppers, R., Hansmann, M.L. & Hartmann, S. JUNB, DUSP2, SGK1, SOCS1 and CREBBP are frequently mutated in T-cell/histiocyte-rich large B-cell lymphoma. Haematologica 104, 330–337 (2019).PMC6355500.

52. Cheng, H., Liu, P., Wang, Z.C., Zou, L., Santiago, S., Garbitt, V., Gjoerup, O.V., Iglehart, J.D., Miron, A., Richardson, A.L., Hahn, W.C. & Zhao, J.J. SIK1 couples LKB1 to p53-dependent anoikis and suppresses metastasis. Science signaling 2, ra35 (2009).PMC2752275.

53. Wood, B., Sikdar, S., Choi, S.J., Virk, S., Alhejaily, A., Baetz, T. & LeBrun, D.P. Abundant expression of interleukin-21 receptor in follicular lymphoma cells is associated with more aggressive disease. Leukemia & lymphoma 54, 1212–1220 (2013).

54. Bhatt, S., Matthews, J., Parvin, S., Sarosiek, K.A., Zhao, D., Jiang, X., Isik, E., Letai, A. & Lossos, I.S. Direct and immune-mediated cytotoxicity of interleukin-21 contributes to antitumor effects in mantle cell lymphoma. Blood 126, 1555–1564 (2015).PMC4582332.

55. Liu, Y., Hu, S., Onder, O., Sahasrabuddhe, A., Rajabi, A., Seitz, A., de Jong, D., Koerts, J., Visser, L., Dzikiewicz-Krawczyk, A., Lim, M.S., Elenitoba-Johnson, K.S., van den Berg, A., Ziel-Swier, L. & Kluiver, J. Palmitoylation by ZDHHC family members regulate B-cell lymphoma growth. Int J Biol Macromol 320, 145876 (2025).

56. Schwaenen, C., Viardot, A., Berger, H., Barth, T.F., Bentink, S., Dohner, H., Enz, M., Feller, A.C., Hansmann, M.L., Hummel, M., Kestler, H.A., Klapper, W., Kreuz, M., Lenze, D., Loeffler, M., Moller, P., Muller-Hermelink, H.K., Ott, G., Rosolowski, M., Rosenwald, A., Ruf, S., Siebert, R., Spang, R., Stein, H., Truemper, L., Lichter, P., Bentz, M., Wessendorf, S. & Molecular Mechanisms in Malignant Lymphomas Network Project of the Deutsche, K. Microarray-based genomic profiling reveals novel genomic aberrations in follicular lymphoma which associate with patient survival and gene expression status. Genes Chromosomes Cancer 48, 39–54 (2009).

57. Mandelbaum, J., Bhagat, G., Tang, H., Mo, T., Brahmachary, M., Shen, Q., Chadburn, A., Rajewsky, K., Tarakhovsky, A., Pasqualucci, L. & Dalla-Favera, R. BLIMP1 is a tumor suppressor gene frequently disrupted in activated B cell-like diffuse large B cell lymphoma. Cancer Cell 18, 568–579 (2010).PMC3030476.

58. Boi, M., Zucca, E., Inghirami, G. & Bertoni, F. PRDM1/BLIMP1: a tumor suppressor gene in B and T cell lymphomas. Leuk Lymphoma 56, 1223–1228 (2015).

59. Bouzelfen, A., Alcantara, M., Kora, H., Picquenot, J.M., Bertrand, P., Cornic, M., Mareschal, S., Bohers, E., Maingonnat, C., Ruminy, P., Adriouch, S., Boyer, O., Dubois, S., Bastard, C., Tilly, H., Latouche, J.B. & Jardin, F. HACE1 is a putative tumor suppressor gene in B-cell lymphomagenesis and is down-regulated by both deletion and epigenetic alterations. Leuk Res 45, 90–100 (2016).

60. Wenzl, K., Manske, M.K., Sarangi, V., Asmann, Y.W., Greipp, P.T., Schoon, H.R., Braggio, E., Maurer, M.J., Feldman, A.L., Witzig, T.E., Slager, S.L., Ansell, S.M., Cerhan, J.R. & Novak, A.J. Loss of TNFAIP3 enhances MYD88(L265P)-driven signaling in non-Hodgkin lymphoma. Blood Cancer J 8, 97 (2018).PMC6177394.

61. Oricchio, E., Nanjangud, G., Wolfe, A.L., Schatz, J.H., Mavrakis, K.J., Jiang, M., Liu, X., Bruno, J., Heguy, A., Olshen, A.B., Socci, N.D., Teruya-Feldstein, J., Weis-Garcia, F., Tam, W., Shaknovich, R., Melnick, A., Himanen, J.P., Chaganti, R.S. & Wendel, H.G. The Eph-receptor A7 is a soluble tumor suppressor for follicular lymphoma. Cell 147, 554–564 (2011).PMC3208379.

62. Xue, W., Kitzing, T., Roessler, S., Zuber, J., Krasnitz, A., Schultz, N., Revill, K., Weissmueller, S., Rappaport, A.R., Simon, J., Zhang, J., Luo, W., Hicks, J., Zender, L., Wang, X.W., Powers, S., Wigler, M. & Lowe, S.W. A cluster of cooperating tumor-suppressor gene candidates in chromosomal deletions. Proc Natl Acad Sci U S A 109, 8212–8217 (2012).PMC3361457.

63. Lang, S., Nguyen, D., Bhadra, P., Jung, M., Helms, V. & Zimmermann, R. Signal Peptide Features Determining the Substrate Specificities of Targeting and Translocation Components in Human ER Protein Import. Front Physiol 13, 833540 (2022).PMC9309488.

64. Scheich, S., Chen, J., Liu, J., Schnutgen, F., Enssle, J.C., Ceribelli, M., Thomas, C.J., Choi, J., Morris, V., Hsiao, T., Nguyen, H., Wang, B., Bolomsky, A., Phelan, J.D., Corcoran, S., Urlaub, H., Young, R.M., Haupl, B., Wright, G.W., Huang, D.W., Ji, Y., Yu, X., Xu, W., Yang, Y., Zhao, H., Muppidi, J., Pan, K.T., Oellerich, T. & Staudt, L.M. Targeting N-linked Glycosylation for the Therapy of Aggressive Lymphomas. Cancer Discov 13, 1862–1883 (2023).PMC10524254.

65. Zambon, A.C., Zhang, L., Minovitsky, S., Kanter, J.R., Prabhakar, S., Salomonis, N., Vranizan, K., Dubchak, I., Conklin, B.R. & Insel, P.A. Gene expression patterns define key transcriptional events in cell-cycle regulation by cAMP and protein kinase A. Proc Natl Acad Sci U S A 102, 8561–8566 (2005).PMC1150853.

66. Grau, M., Lopez, C., Martin-Subero, J.I. & Bea, S. Cytogenomics of B-cell non-Hodgkin lymphomas: The “old” meets the “new”. Best Pract Res Clin Haematol 36, 101513 (2023).

67. Dias, L.M., Thodima, V., Friedman, J., Ma, C., Guttapalli, A., Mendiratta, G., Siddiqi, I.N., Syrbu, S., Chaganti, R.S. & Houldsworth, J. Cross-platform assessment of genomic imbalance confirms the clinical relevance of genomic complexity and reveals loci with potential pathogenic roles in diffuse large B-cell lymphoma. Leukemia & lymphoma 57, 899–908 (2016).PMC4963821.

68. Wu, W., Lu, P., Patel, P., Ma, J., Cai, K.Q., Mallikarjuna, V.S., Poureghbali, S., Nakhoda, S.R., Nejati, R. & Lynn Wang, Y. SHP1 loss augments DLBCL cellular response to ibrutinib: a candidate predictive biomarker. Oncogene 42, 409–420 (2023).

69. Bott, A.J., Shen, J., Tonelli, C., Zhan, L., Sivaram, N., Jiang, Y.P., Yu, X., Bhatt, V., Chiles, E., Zhong, H., Maimouni, S., Dai, W., Velasquez, S., Pan, J.A., Muthalagu, N., Morton, J., Anthony, T.G., Feng, H., Lamers, W.H., Murphy, D.J., Guo, J.Y., Jin, J., Crawford, H.C., Zhang, L., White, E., Lin, R.Z., Su, X., Tuveson, D.A. & Zong, W.X. Glutamine Anabolism Plays a Critical Role in Pancreatic Cancer by Coupling Carbon and Nitrogen Metabolism. Cell Rep 29, 1287–1298 e1286 (2019).PMC6886125.

70. Werner, A., Schafer, S., Gleussner, N., Nimmerjahn, F. & Winkler, T.H. Determining immunoglobulin-specific B cell receptor repertoire of murine splenocytes by next-generation sequencing. STAR Protoc 3, 101277 (2022).PMC9010798.

71. Jung, J., Zhu, S., Lalani, A., Shakarchi, J., Matracz, B., Wu, G.G., Zong, W.X., Zhao, L. & Xie, P. Commensal Bacteria Drive B-cell Lymphomagenesis in the Setting of Innate Immunodeficiency. Blood Cancer Discov 6, 505–525 (2025).PMC12279395.

